# *In situ* structural insights into the excitation contraction coupling mechanism of skeletal muscle

**DOI:** 10.1101/2023.08.26.554922

**Authors:** Jiashu Xu, Chenyi Liao, Chang-Cheng Yin, Guohui Li, Yun Zhu, Fei Sun

**Author notes:** Correspondences: Guohui Li, Yun Zhu or Fei Sun. First authors with equal contributions (Jiashu Xu, Chenyi Liao).

## Abstract

Excitation–contraction coupling (ECC) is a fundamental mechanism in control of skeletal muscle contraction and occurs at triad junctions, where dihydropyridine receptors (DHPRs) on transverse tubules sense excitation signals and then cause calcium release from the sarcoplasmic reticulum via coupling to type 1 ryanodine receptors (RyR1s), inducing the subsequent contraction of muscle filaments. However, the molecular mechanism remains unclear due to the lack of structural details. Here, we explored the nanometre-resolution architecture of triad junction by cryo-electron tomography, solved the *in situ* structure of RyR1 in complex with FKBP12 and calmodulin, and discovered the intact RyR1-DHPR supercomplex. RyR1s arrange into two rows on the terminal cisternae membrane by forming right-hand corner-to-corner contacts, and tetrads of DHPRs bind to RyR1s in an alternating manner, forming another two rows on the transverse tubule membrane. Such unique arrangement is important for synergistic calcium release and provides direct evidence of physical coupling in ECC.

## Introduction

Muscle contractions enable essential animal behaviours, such as cardiac pulsations, exercise, respiration and maintenance of posture. Excitation-contraction coupling (ECC) is a fundamental process that links the action potential to the contraction of striated muscle fibres in vertebrates (*1*). Upon arrival of action potentials at the neuromuscular junction in skeletal musculature, calcium inflow into presynaptic terminals triggers exocytosis of acetylcholine (ACh)-containing sacs into synaptic clefts, leading to depolarization of the plasma membrane. The depolarization propagates along transverse tubules (T-tubules), which are invaginations of the sarcolemma that penetrate deep into skeletal muscle fibres (*2, 3*). Triad junctions are formed in skeletal muscle by T-tubules sandwiches between the two sides of enlarged portions of the sarcoplasmic reticulum (SR), named terminal cisternae (TC), ensuring an efficient ECC process (*4*). Then, in triad junctions, voltage-sensitive dihydropyridine receptors (DHPRs) in the T-tubule membrane (TTM) interact with type 1 ryanodine receptors (RyR1s) in the adjacent TC membrane (TCM), leading to a rapid release of Ca^2+^ from the SR and the subsequent contraction of myofibrils (*5–8*).

As the high-conductance Ca^2+^ channel localized at the SR, RyR is the largest known ion channel, with ∼5,000 residues in each protomer and a homotetrameric organization of over 2.2 million Daltons (MDa) (*9, 10*). It consists of a relatively small transmembrane region forming an ion-conducting pore and a large cytoplasmic region responsible for interacting with various ligands to regulate the channel (*11*). Mammals have three RyR isoforms, named RyR1, RyR2 and RyR3. RyR1 and RyR2 are expressed at high levels in skeletal and cardiac muscles, respectively (*12–14*). In triad junctions of skeletal muscle, RyR1s are anchored to TCM in close proximity, thereby constituting efficient Ca^2+^ release units (CRU) with DHPRs located on the T-tubules. Upon arrival of the action potential, DHPR triggers opening of the RyR1 channel through mechanical coupling, but the details of this process remain enigmatic.

In the 1980s, high-density ‘foot’ structures were discovered in the triad junctions of frog twitch fibres (*15*). Further investigation demonstrated that these structures were RyR1 channels (*16*). In 1989, RyR1 was purified from the fast-twitch skeletal muscle in rabbit, and its distinctive mushroom-shaped structure was resolved at 3.7 nm resolution (*17*). With the rapid development of cryo-electron microscopy (cryo-EM) in recent years, numerous high-resolution structural studies from purified specimens have further illuminated different states of RyR1 channels (*18–22*). However, how RyR1s are activated and regulated *in situ* in skeletal muscle is unclear.

At the triad junctions of skeletal muscle, RyR1 typically appears *in situ* in two (sometimes three) parallel rows (*23–25*). This pattern is conserved across crustaceans and vertebrates (*23–26*). In swim bladder muscle fibres from toadfish, adjacent RyR1 channels were found to be arranged in a slightly twisted corner-to-corner manner (*27*). Other studies revealed that when crystallized into two-dimensional (2D) arrays, RyR1 channels form a checkerboard-like pattern with adjacent pairs bound together in a similar corner-to-corner way (*28*).

In addition to the arrangement of RyR1s in the triad junctions, how RyR1s interact with DHPRs at the triad junctions remains a topic of great research interest because it is related to the molecular mechanism of ECC. Previous studies using the freeze-fracture technique revealed diamond-shaped particle clusters named “tetrads” on TTMs, suggesting a direct mechanical interaction between RyR1 and DHPR (*29, 30*). Although different possible interaction sites and modes between RyR1 and DHPR have been proposed in recent decades (*31–41*), direct evidence is still missing to confirm the existence of their physical interactions.

Recent trails of *in situ* structural studies using cryo-electron tomography (cryo-ET) revealed the presence of RyR1-like density on the TCM of triad junctions from cryo-lamellae of toadfish swim-bladder muscle prepared by the cryo-focused ion beam (cryo-FIB) technique (*42*). Besides, SR vesicles purified from rabbit skeletal muscle were used to solve the *in situ* structure of RyR1 at nanometre resolution (*43*), and the resolution was updated to subnanometer 9.1 Å in recent years (*44, 45*). However, in these studies, no DHPRs were observed on the TTM due to the low resolution of the tomogram (*42*) or to potential specimen damage during the purification procedure (*43, 44*). We recently developed a workflow, VHUT-cryo-FIB, for cryo-lamellae preparation from tissue specimens and successfully observed clear densities within the triad junctions of rat skeletal muscle in the tomogram (*46*), providing a potential method to obtain high-resolution *in situ* structures of RyR1/DHPR in skeletal muscle.

In this study, we explored another specimen preparation procedure to fabricate cryo-lamellae of mouse skeletal muscle using cryo-FIB and then performed an *in situ* cryo-ET study of intact triad junctions. Both RyR1 and DHPR densities were clearly observed within the triad junctions in our tomograms. Then, we resolved the *in situ* structures of both the RyR1 tetramer and RyR1-DHPR supercomplex with nanometre resolution using subtomogram averaging (STA). We further deduced the ordered spatial arrangements of RyR1s and DHPRs within the triad junctions, which were further investigated by molecular dynamics simulations, suggesting the mechanism of synergic coupling between RyR1 and DHPR and among RyR1s. Our results provide deep structural insights into the physiology of triad junctions of mammalian skeletal muscle and highlight new avenues towards better understanding the ECC mechanism in skeletal muscle.

## Results

### Overall intact structure of triad junction

To preserve the native integrity of triad junctions, we prepared cryo-lamellae from mouse skeletal muscle tissue by improving the previous protocol (*47*). Specifically, we separated intact skeletal muscle fibres from the extensor digitorum longus (EDL) muscle of 2∼3-week-old mice (**Figs. 1a and S1a**). The fibres were exposed to a 10% glycerol solution for several minutes before placing onto a cryo-EM grid, which ensured complete vitrification during subsequent plunge freezing (*48–50*). Then, the vitrified specimen was milled to a thickness of approximately 150 nm using cryo-FIB, and the cryo-lamellae were subjected to cryo-ET data collection by targeting the triad junctions manually (**Figs. S1b-e and Table S1**). The textures of muscle actin filaments were prominently visible in the raw cryo-EM micrograph, and numerous strips constituting the triad junction could be found perpendicular to the filaments and aligned parallelly and orderly (**Fig. 1b**). We measured the distances between two adjacent parallel triad junctions and found that they ranged from 594 nm to 1,171 nm with a mean of 833 nm and a standard deviation of 122 nm (**Fig. 1c**).

**Figure 1.**
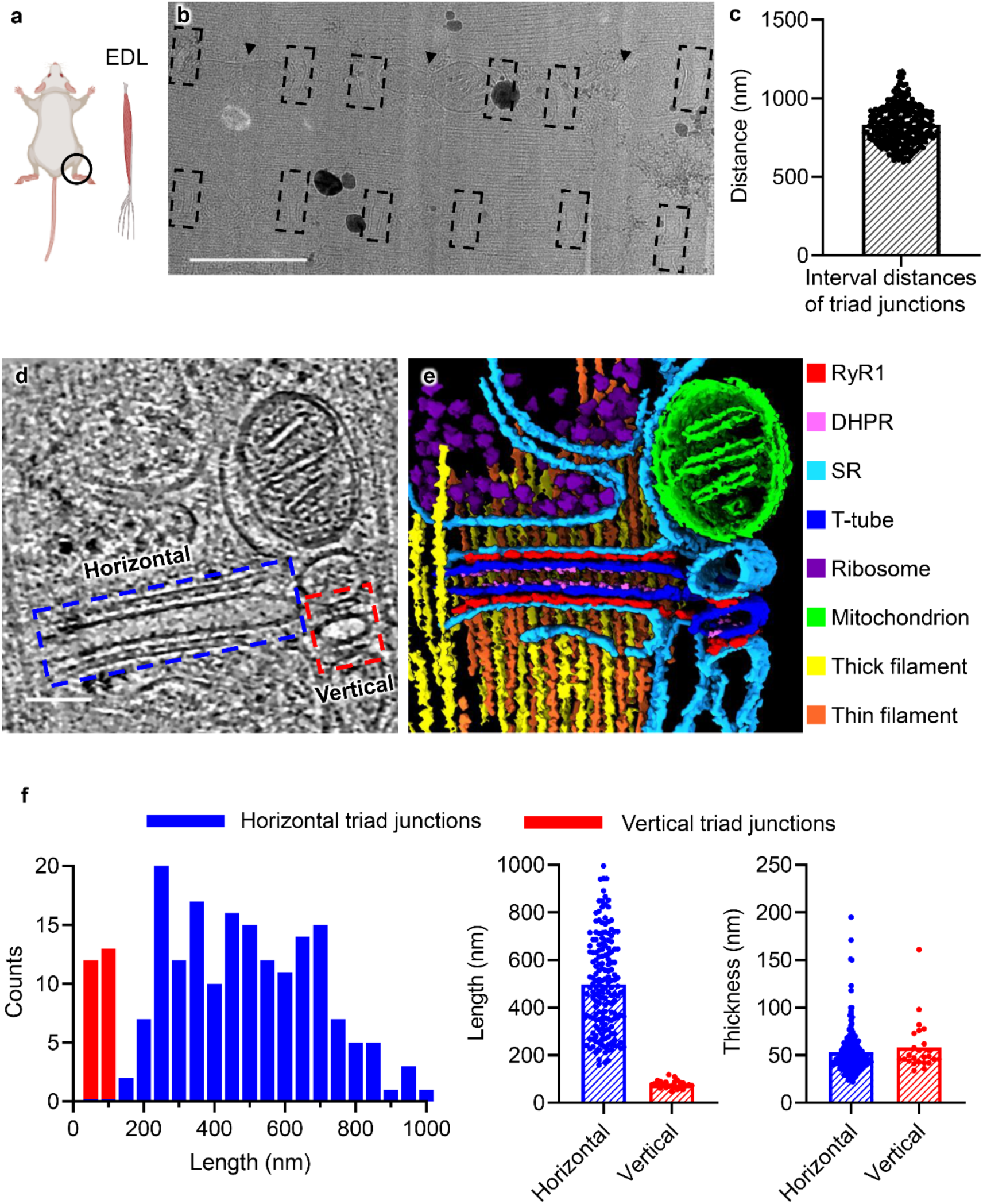
Overall architecture of native triad junctions in mouse skeletal muscle. (**a**) EDL muscle fibres were manually extracted from the mouse leg, and the black circle indicates the extraction position. (**b**) Low magnification cryo-EM image of cryo-lamella of mouse skeletal muscle. Regularly arranged triad junctions are marked by black dotted line boxes. The mitochondria near triad junctions are marked by black triangles. Scale bar, 1 μm. (**c**) The distribution of the interval distances among the triad junctions. (**d**) A representative tomogram slice shows two perpendicular triad junctions and their surrounding environment. Scale bar, 100 nm. Horizontal and vertical triad junctions are marked by blue and red dotted line boxes, respectively. (**e**) The segmented tomogram shows different components around triad junctions in (d), including RyR1 (red), DHPR (pink), SR (cyan), T-tubule (blue), ribosome (purple), mitochondrion (green), thick filament (yellow), and thin filament (orange). The segmentation was performed by using the Microscopy Image Browser (MIB) tool (*101*). (**f**) The distributions of length/width and thicknesses of T-tubules in horizontal (blue) and vertical (red) triad junctions.

From a representative tomogram (**Figs. 1d-e and Movie S1**), we observed useful information about the ultrastructure around the triad junctions. The T-tubule is enclosed by TCM on both sides with RyR1s clearly on the TCM and putative DHPRs on the TTM. Thick and thin filaments dominate most of the cellular space and are proximal to triad junctions, ready to perceive calcium signals from SR and then trigger dynamic sliding of myosin on actin filaments. Mitochondria were found to be strategically localized near the triad junctions, where they can offer critical energy necessary for the sliding of myosin. In addition, we found abundant ribosome particles surrounding the SR, which are important for the protein metabolism of both triad junctions and myofilaments. The presence of two perpendicular triad junctions discloses the narrow and flattened shape of T-tubules, which form a network inside the skeletal muscle and attach to the flat and elongated surfaces of TCs.

We then performed statistical analyses of the sizes of T-tubules by using a total of 198 tomograms containing the most complete triad junctions (**Fig. 1f**). The measured apparent lengths of T-tubules vary continuously from 49 nm to 996 nm because of the randomly distributed angles between the long axes of T-tubules and the direction of cryo-FIB milling. The dual-modular distribution of the apparent lengths indicates two categories of T-tubules, with one having an apparent length ranging from 160∼996 nm, representing the tilted and horizontal T-tubules, and the other having an apparent length ranging from 49∼118 nm, representing the close-to-vertical T-tubules. For the vertical T-tubules, there are only two RyR1 molecules located on TCMs, while for the tilt-and- horizontal T-tubules, there are more RyR1s observed.

Notably, the measured apparent lengths of vertical T-tubules represent their widths. Thus, we were able to calculate the average length of T-tubules as 498 nm with a standard deviation of 197 nm and the average width of T-tubules as 77 nm with a standard deviation of 16 nm (**Fig. 1f**). Due to the limited imaging area of each tomogram with a limited coverage of triad junctions, we believe there would be many triad junctions much longer in length than they appear in tomograms. It is noteworthy that the thicknesses of both horizontal and vertical T-tubules are almost the same, with mean values of 53 nm and 58 nm, respectively (**Fig. 1f**), suggesting potential physiological relevance.

### *In situ* structure of RyR1 in native triad junctions

We then manually picked out the foot-like particles that represent RyR1s located between the TTM and TCM and performed subtomogram analysis (**Fig. S2a**), resulting in a final averaged cryo-EM map of *in situ* RyR1 (**Fig. 2a, Table S1 and Movie S2**) with a resolution of 16.7 Å based on the gold standard criteria of Fourier shell correlation (**Fig. S2c**). In addition to the typical square shape of RyR1 in its top view with a width of 26.7 nm, we observed both TCM and TTM in the cryo-EM map. The transmembrane region of RyR1 is embedded in the TCM, and the distance between the TCM and TTM can be measured as 16.7 nm, which leaves space for the interaction between RyR1 and DHPR.

**Figure 2.**
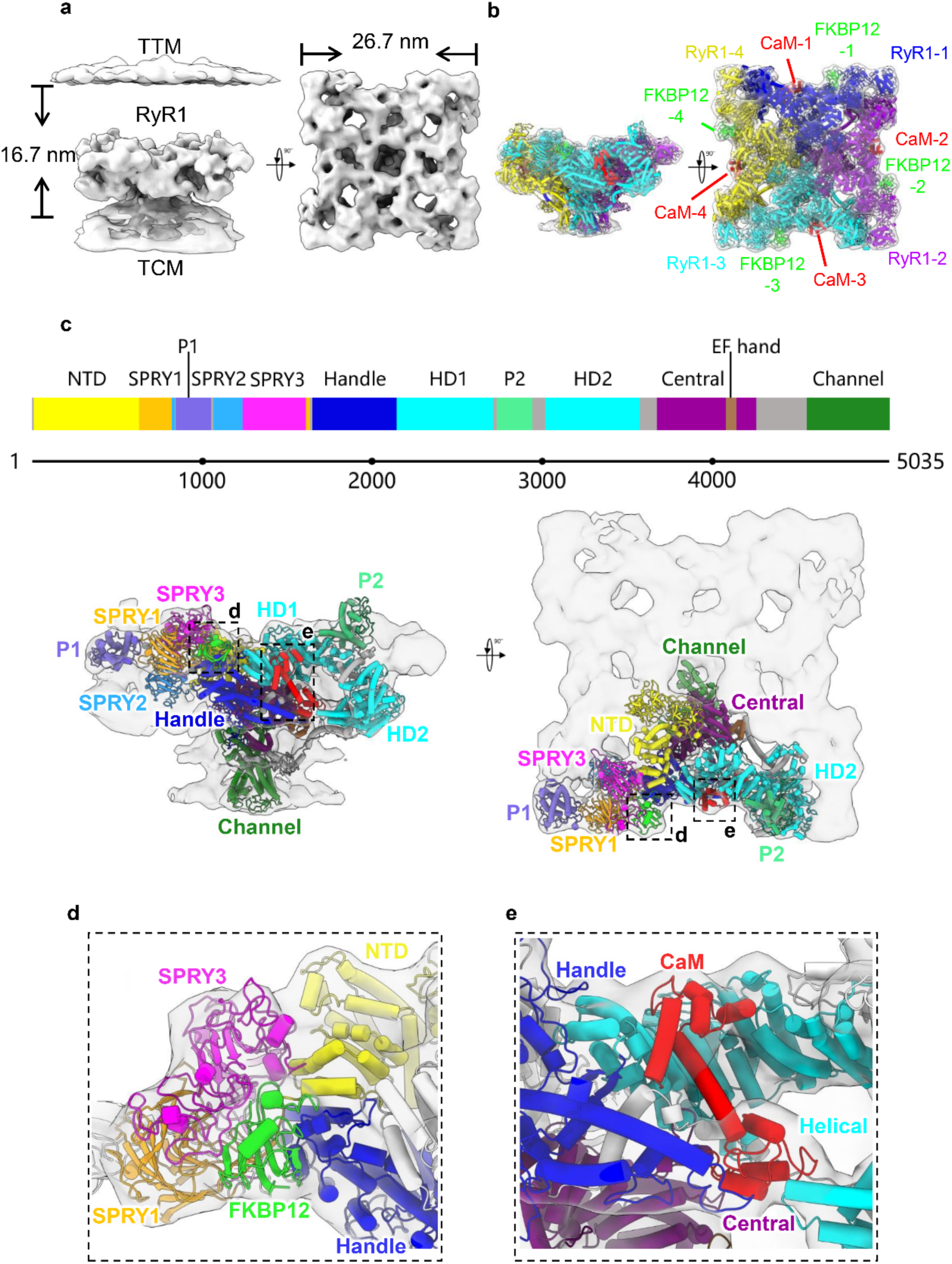
*In situ* structure of RyR1 in the native triad junction. (**a**) The side and top views of cryo-EM map of *in situ* RyR1 embedded in TCM and proximal to TTM. (**b**) The structural model of the RyR1-FKBP12-apo-CaM complex was flexibly fitted into the cryo-EM map in (a). FKBP12s, apo-CaMs and four RyR1 protomers are indicated accordingly. (**c**) Distribution and definition of mouse RyR1 domains with one protomer fitted into the cryo-EM map. Locations of FKBP12 and apo-CaM are indicated by the black dotted boxes. (**d**) Zoomed-in view of the FKBP12 region in (c). (**e**) Zoomed-in view of the apo-CaM region in (c).

The cryo-EM map of *in situ* RyR1 can be well fitted with the previously reported high-resolution structures of RyR1s in both open (PDB entry 5J8V, **Fig. S3a**) (*22*) and closed (PDB entry 4UWA, **Fig. S3b**) (*18*) states, which could not be distinguished at the current resolution. However, we also observed several different pieces of densities, one piece around the clamp region and other pieces around the handle and helical domains, which would represent the binding partners of RyR1, including FK506 binding protein 12 (FKBP12) and calmodulin (CaM), respectively.

FKBP12, which is alternatively known as calstabin1 and abundantly expressed in skeletal muscle, has been reported to interact physically with RyR1 and effectively stabilize the closed state of RyR1 (*51, 52*). The high-resolution structure of the RyR1-FKBP12 complex (PDB entries 3J8H and 5TB0) (*19, 21*) fits appropriately into the cryo-EM map of *in situ* RyR1, leaving distinct pieces of densities around the handle and helical domains, which would represent the density of CaM (**Figs. S3c-d**).

CaM is another key binding partner that can regulate the activity of RyRs, while in skeletal muscle, the effect of this regulation varies depending on the concentration of Ca^2+^ (*53*). At nanomolar concentrations of Ca^2+^, CaM is in its apo form (apo-CaM), which can bind to RyR1 and weakly activate the opening of RyR1, while at micromolar concentrations of Ca^2+^, CaM is in its calcium binding form (Ca^2+^-CaM), which will bind to RyR1 with a different binding site and inhibit the opening of RyR1 (*52, 54, 55*). In recent years, the high-resolution structures of RyR1 bound with CaM were determined with (PDB entry 7TZC) and without Ca^2+^ (PDB entry 6X33) in the buffer (*56, 57*). However, both structures were supposed to be closer to the apo-CaM bound state (RyR1-apo-CaM) because the binding position of CaM in the structure of Ca^2+^ loading RyR2-CaM complexes (*58*) shows a slight slipping down compared to these two structures. These two structures can be fitted into our cryo-EM map of *in situ* RyR1, including the abovementioned pieces of densities (**Figs. S3e-f**). In addition, we found that the density near the HD1 domain in our cryo-EM map is absent in the low-pass filtered map of RyR2-Ca^2+^-CaM (EMDB entry EMD-9833) but exists in the low-pass filtered map of RyR2-apo-CaM (EMDB entry EMD-9836) (*58*) (**Fig. S3g**). These structural analyses suggest that the present *in situ* structure of RyR1 represents the apo-CaM binding state of RyR1, which is further stabilized by the binding of FKBP12. Considering that we did not trigger myofibrils with additional calcium buffer during specimen preparation, we speculate that the present *in situ* structure of RyR1 represents a closed steady state.

Based on a combination of homology modelling (**Fig. S4**) and artificial intelligence modelling, we built a full-length model of mouse RyR1 that was fitted into the *in situ* cryo-EM map together with the fitting of both FKBP12 and apo-CaM. Then, the molecular dynamics flexible fitting (MDFF) approach was utilized to obtain an integrative *in situ* structural model of the RyR1-FKBP12-apo-CaM complex (**Figs. 2b-c and Movie S2**). In this model, FKBP12 is encapsulated by the N-terminal domain (NTD), SPRY1 and 3 domains and handle domain (**Fig. 2d**), while apo-CaM is surrounded by the central, handle and helical domains (**Fig. 2e**). Notably, the connection between apo-CaM and the HD2 domain at the bottom of the clamp region could only be found in the present *in situ* map (**Fig. S3h**) and was missing in the previously reported cryo-EM map of the RyR1-apo-CaM complex (*56, 57*), indicating a stabilized state of *in situ* RyR1 in the present study.

### Arrangement of RyR1s in native triad junctions

To investigate the *in situ* organization of RyR1s, we mapped the aligned particles of RyR1 tetramers back to their original tomograms and found that RyR1 tetramers are predominantly arranged one-by-one in two parallel rows within the triad junctions (**Figs. 3a, S5 and Movie S3**). Moreover, two neighbouring RyR1 tetramers interact with each other in a right-hand corner-to-corner manner (**Figs. 3a and S5**), which means that when viewed from the cytoplasmic side of RyR1, the interaction interfaces of neighbouring RyR1 tetramers are always located at the right side of the corner of each tetramer. Such a right-hand corner-to-corner array was suggested by previous studies of 2D crystals of the RyR1 tetramer (*28*), a thin section study of triad junctions (*59*) and a low-resolution cryo-ET study based on isolated triad junctions (*43*). In this study, we not only resolved the right-hand corner-to-corner arrangement of RyR1 tetramers directly at high resolution but also quantified the distances between two adjacent tetramers, which ranged primarily (85%) from 29 to 33 nm with an average of 31 nm (**Fig. 3b**).

**Figure 3.**
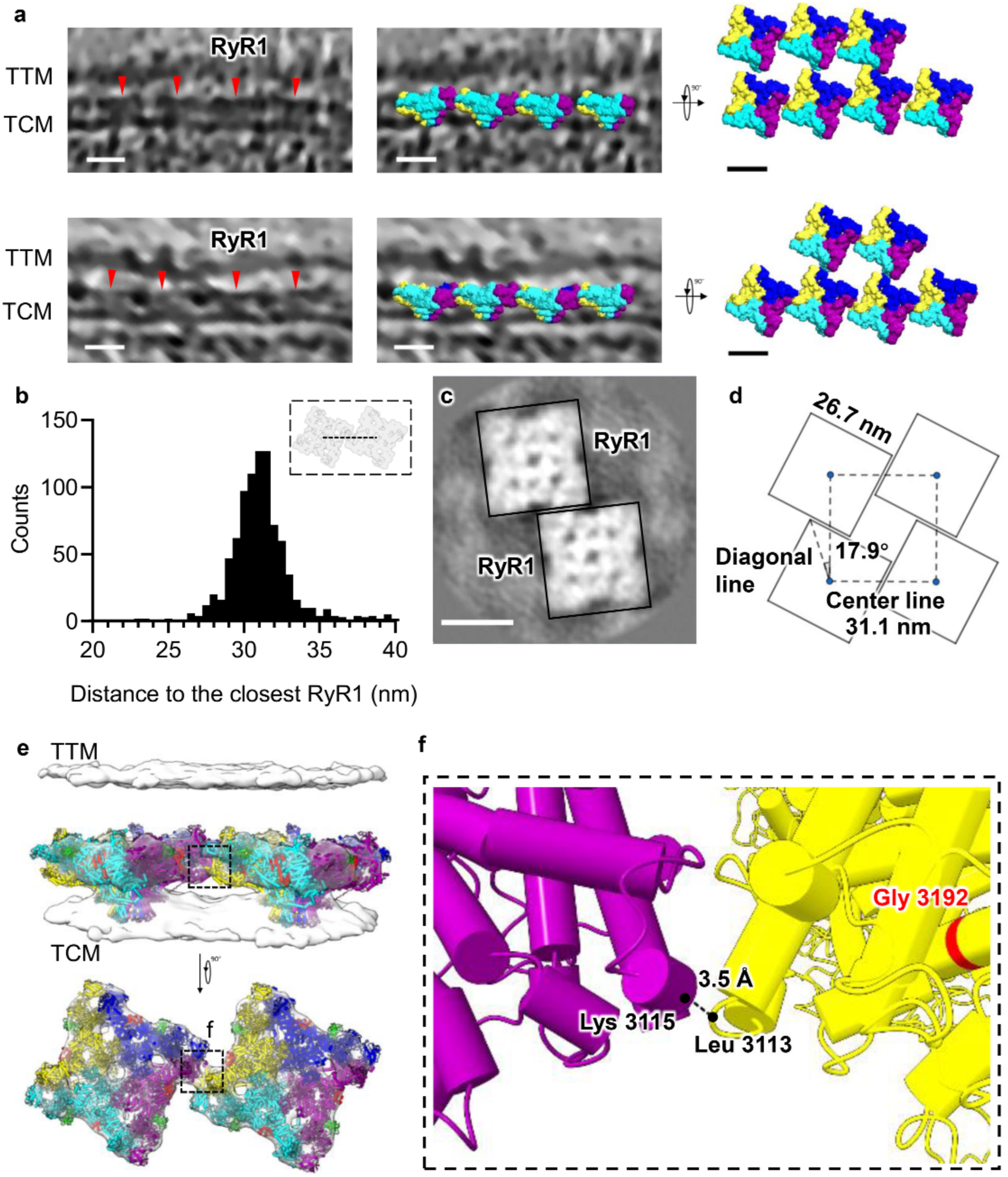
*In situ* arrangement of RyR1s in the native triad junction. (**a**) Two representative tomogram slices showing the embedded array of RyR1 tetramers in the triad junctions. The structures of RyR1s are plotted back into the tomograms according to their refined coordinates and Euler angles. Scale bar, 20 nm. (**b**) The statistical distribution of distances between two neighbouring RyR1 tetramers. (**c**) A slice of the STA averaged map of dimeric RyR1 tetramers. Scale bar, 20 nm. (**d**) Diagram of the geometric arrangement of RyR1 tetramers, which are represented as square boxes. (**e**) The STA averaged map of dimeric RyR1 tetramers in the triad junction, showing both TTM and TCM. The structural model of RyR1-FKBP12-apo-CaM is fitted into the map. The contact site between two tetramers is marked by the black dotted box. (**f**) The zoomed-in view of the contact site in (e). The nearest potential contacting residues Lys3115 and Leu3113 are labelled with their main-chain distance indicated. The position of residue Gly3192, whose mutation was reported to be relevant to premature mortality in two patients (*61*), is indicated in red.

To confirm the exact interaction between two adjacent RyR1 tetramers, we resubtracted the subtomograms of neighbouring tetramers and then performed subtomogram averaging based on the original orientation parameters of a single tetramer, resulting in a clear density at the corner-to-corner interaction site of two neighbouring tetramers (**Fig. 3c, Table S1 and Movie S3**). The centre-to-centre separation between these two tetramers was measured as 31.1 nm (**Fig. 3d**), which is consistent with the above statistics. The length of the square edge of the RyR1 tetramer was also measured as 26.7 nm, and the angle between the diagonal line of the tetramer and the central line of the RyR1 array was measured as 17.9° (**Fig. 3d**), which is related to the exact position of the corner interaction. These precise measurements illustrate the geometric arrangement of RyR1 tetramers in the native triad junction, providing direct proof of previous suggestions from 2D crystallographic studies (*28*).

We then fitted the structural model of the RyR1-FKBP12-apo-CaM complex into the averaged map of dimeric RyR1 tetramers (**Fig. 3e**) and observed the potential contact site of RyR1 tetramers that is situated in the HD2 domain, specifically with residues ranging from 3087 to 3118 (**Fig. 3f**). We found that the nearest site lies between Leu3113 of one RyR1 tetramer and Lys3115 of another tetramer, and their main-chain distance is less than 3.5 Å, suggesting a potential direct interaction at this site (**Fig. 3f**). Indeed, previous studies have suggested that the contact site of neighbouring RyR1 tetramers is located at residues ranging from 2540 to 3207 (*28, 60*). Furthermore, it has been reported that the point mutation of Gly3191 in RyR1, located proximal to the contact site, is causative for premature mortality in two patients (*61*). According to the structural model of the dimeric RyR1 tetramers here, we speculate that the alteration of Gly3191 would strongly and negatively affect the corner-to-corner interaction between neighbouring tetramers and thus eliminate the coupling between them, which will be further studied below.

### Molecular dynamics insight into the coupling of neighbouring RyR1s

The specific array arrangement of RyR1 tetramers with corner-to-corner interactions suggests its essential role in the conformational change coupling of RyR1s in one triad junction, which has been hypothesized previously (*28*) but lacks experimental support. Here, based on the *in situ* structural model of the RyR1 tetramer array, we performed molecular dynamics (MD) simulations (**Tables S2 and S3**) to investigate the structural dynamics of RyR1s in the array.

First, coarse-grained (CG) MD simulations were performed to explore the assembly of four RyR1 tetramers (**Fig. S6a and Movies S4-5**). Starting with RyR1 tetramers more than 20 Å away from each other, the RyR1 tetramers in the open conformation (**Movie S4**) tend to form more right-hand corner-to-corner contacts through intermolecular helical-helical domain interactions than those in the closed conformation (**Movie S5**) during 3-μs CG simulations (**Figs. 4a-b**). The assembled architecture formed by the open RyR1s is consistent with their arrangements in the triad junctions *in situ* (**Fig. 4a**). This observation suggests that RyR1s in the open conformation can spontaneously self-assemble into the right-hand corner-to-corner contact pattern.

**Figure 4.**
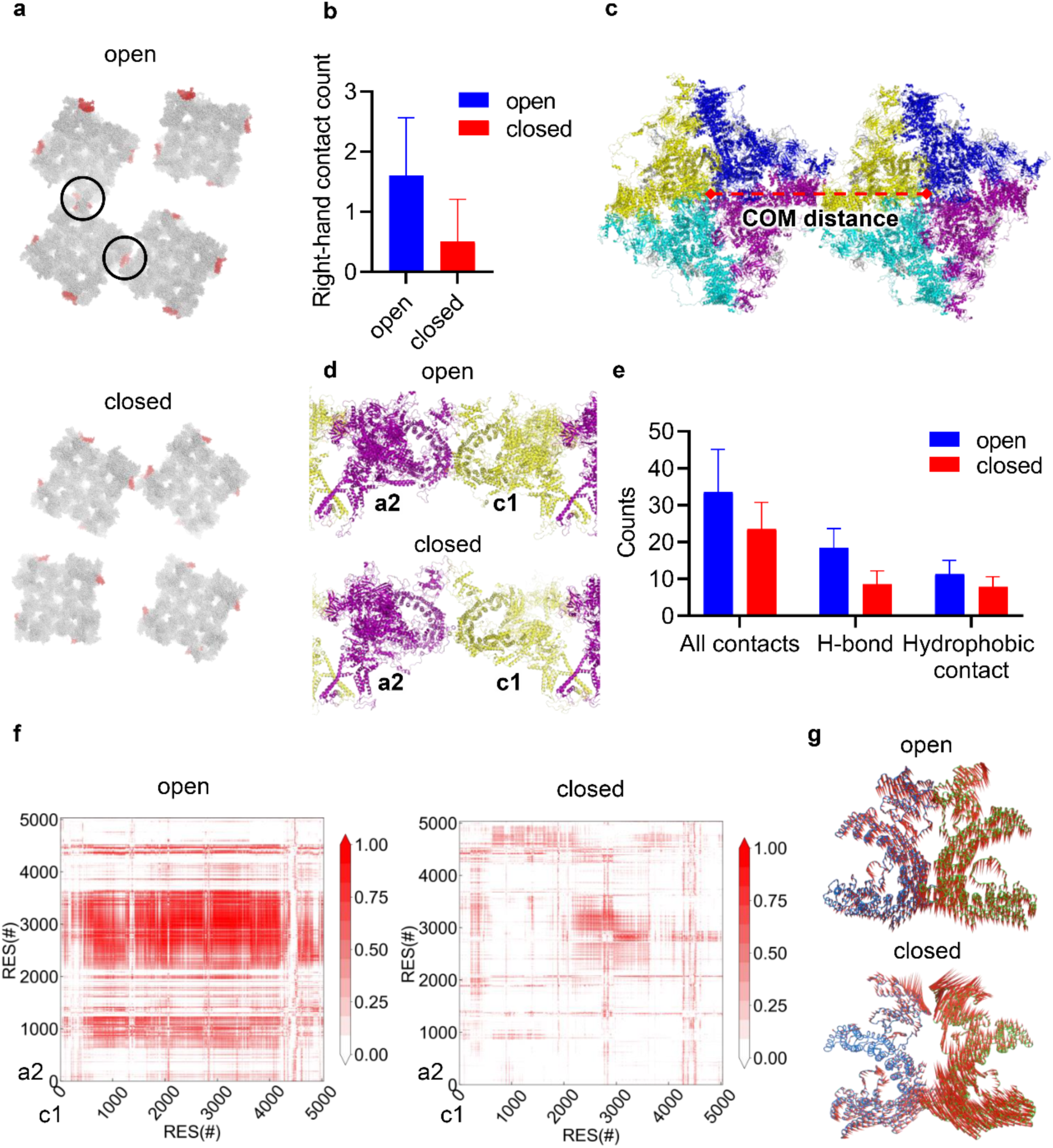
Molecular dynamics (MD) simulations of RyR1-RyR1 interactions and dynamics. (**a**) Representative results of CG MD simulations of four RyR1 tetramers in the open and closed states. Residues ranging from 2951 to 3240 are shown in red, and the right-hand contacts between RyR1s are marked by black circles. (**b**) Statistics of the counts of right-hand corner-to-corner contacts observed during CG MD simulations for the open and closed states. (**c**) Representative dimeric RyR1 tetramers in the open state from all-atom MD simulations. (**d**) Magnified views of two contact RyR1 protomers, a2 in purple and c1 in yellow, in the open and closed states, respectively. (**e**) Statistics of the counts of all contacts (< 6 Å), hydrogen bonds, and hydrophobic contacts (< 6 Å) during the last 110 ns of all-atom MD simulations. (**f**) DCCMs of the interacting RyR1 protomers, a2 and c1 in (d). Correlated values above zero are represented in gradient red, and the high positive value represents a strongly correlated motion. A significant correlative motion can be observed for the open RyR1 protomers. (**g**) The primary motions of residues ranging from 2494 to 3611 at the contact P2 and HD2 domains of the interacting RyR1 protomers in the open and closed states, respectively. These primary motions were derived based on the first principal component (PC1s) from PCA investigation of all-atom MD simulations of dimeric RyR1 tetramers.

Then, we performed all-atom MD simulations to examine the interactions and motions of two adjacent RyR1 tetramers (**Fig. S6b and Movies S6-7**). Starting from the initial separation setup, these two tetramers, in both open and closed conformations, tended to quickly interact with each other through the intermolecular helical-helical domains, and the distance of their centre-of-mass (COM) decreased from ∼330 Å to 315-325 Å in three independent MD replicas (**Figs. 4c and S7a**). However, we observed that the open RyR1s exhibited more intermolecular hydrogen bonds (H-bonds) and hydrophobic contacts than the closed state, resulting in tighter helical-helical domain interaction (**Figs. 4d-e and S7b**). In addition, the open RyR1s showed a stronger dynamic correlation than the closed state (**Figs. 4f and S7c**). Specifically, compared to the closed RyR1s, significant correlative motions between open RyR1s were observed, particularly at the P2 and HD2 domains, with residues ranging from 2200 to 3611. Interestingly, the two most distant RyR1 protomers also showed strong correlative motions in the open state compared to the closed state (**Figs. S7d-e**).

Furthermore, we utilized principal component analysis (PCA) to conduct another comprehensive investigation of the global motions of assembled RyR1s in both open and closed states (**Fig. S8**). We found that the two open RyR1s exhibit a concerted and synergetic motion of up and down swinging at the P2 and helical domains upon contact with each other (**Figs. 4g and S8**). In contrast, such strong correlative and symmetrical movements were not observed for the RyR1s in the closed state (**Figs. 4g and S8**). These analyses indicate that the strong interactions between neighbouring RyR1s in the open state promote the synergetic movement of their P2 and helical domains, providing the structural basis of coupling between neighbouring RyR1s for coactivation and synergistic calcium release, which was previously termed the calcium spark in cardiac muscle (*62*).

### RyR1 interacts with DHPR directly in the native triad junction

DHPR is commonly known as the voltage-gated L-type Ca^2+^ channel, also referred to as Cav1. It comprises several subunits, including the core pore-forming and voltage-sensing α1 subunit, the transmembrane γ subunit, the extracellular α2δ subunit, and the intracellular β1a subunit (*63*). The α1 and β1a subunits have been identified as critical components in the ECC process (*64*). Unlike cardiac muscle, which relies on a “calcium-induced calcium release” mechanism (*65*), DHPRs appear to transmit signals to RyR1s through mechanical coupling in skeletal muscle. Despite extensive research over recent decades in search of evidence for the interaction between RyR1 and DHPR, direct evidence is still absent due to a lack of detectable DHPR densities either in isolated T-tubule vesicles or in fixed skeletal muscle tissues (*42–44*).

In our tomograms featuring native triad junctions, we observed protrusion densities on the extracellular side of TTMs (**Fig. 5a**). These densities exhibited a symmetrical arrangement and were accompanied by RyR1 density on the corresponding TCM, some of which possessed an intracellular portion linked to the RyR1 corner (**Fig. 5b**). Considering previous results (*29*), it is highly likely that these densities anchored to the TTM belong to DHPRs. Intriguingly, many RyR1s lack such accompanying density, suggesting that the copy number of tetrads formed by DHPRs is lower than that of RyR1s in triad junctions (**Fig. 5c**).

**Figure 5.**
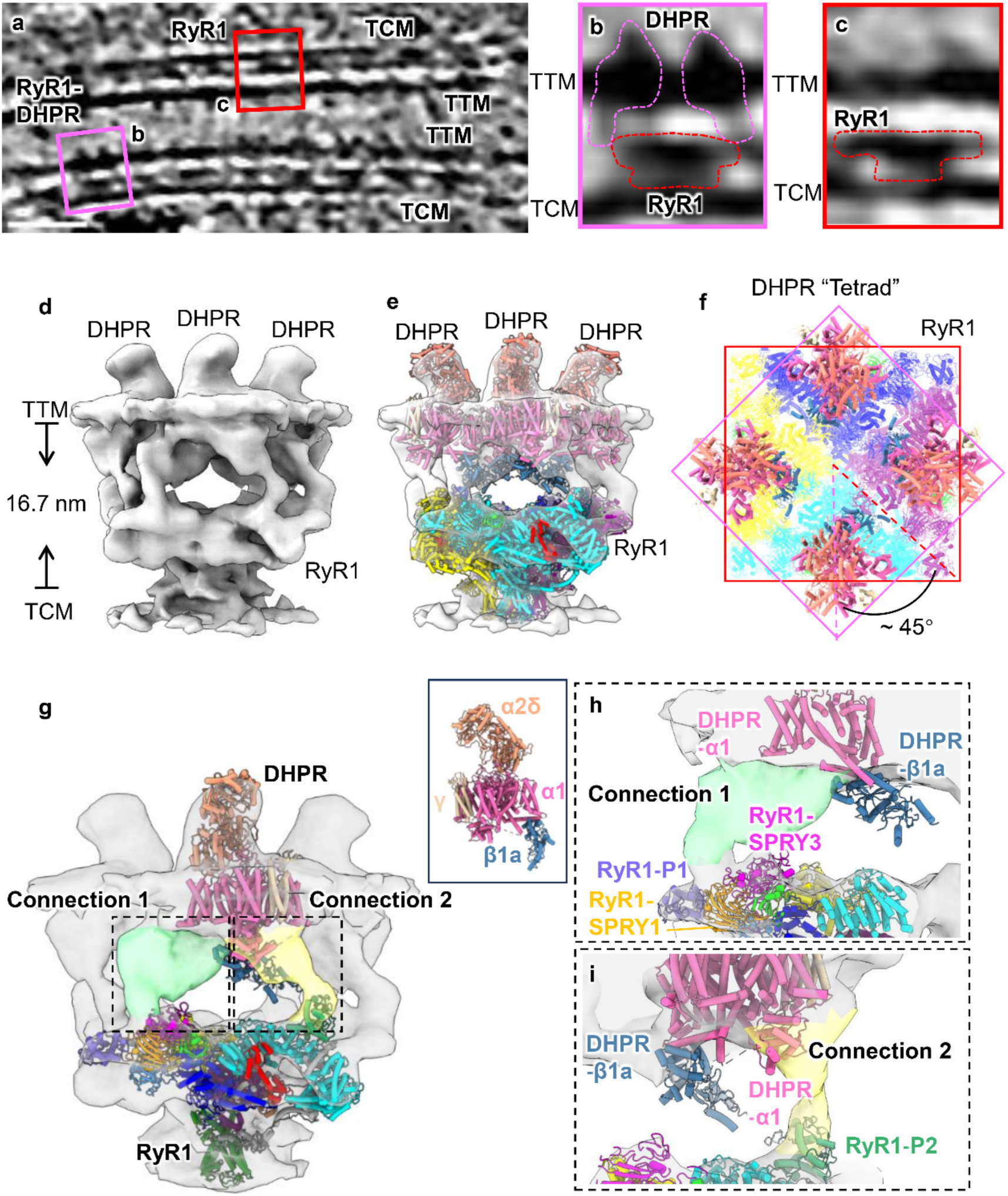
The *in situ* structure of the RyR1-DHPR supercomplex in the native triad junction. (**a**) The representative tomogram slice shows that both RyR1 and DHPR densities existed in the triad junction. The pink box marks a RyR1-DHPR supercomplex, and the red box marks RyR1 without DHPR bound. Scale bar, 100 nm. (**b**) Magnified view of the RyR1-DHPR supercomplex in (a). (**c**) Magnified view of RyR1 without DHPR bound in (a). (**d**) The final averaged map of the RyR1-DHPR supercomplex with TTM and TCM. The distance between TTM and TCM is indicated. (**e**) The structural model of RyR1-FKBP12-apo-CaM and the previously reported structural model of DHPR (PDB entry 5GJW) (*63*) are fitted into the RyR1-DHPR supercomplex map. (**f**) Top view of the RyR1-DHPR supercomplex, showing the twist angle between the “tetrad” formed by four DHPRs and the RyR1 tetramer. (**g**) The two connection densities between RyR1 and DHPR are shown in green and yellow. The RyR1 domains and DHPR components are coloured differently. (**h**) Magnified view of the density map around Connection 1 in (g). (**i**) Magnified view of the density map around Connection 2 in (g).

We manually picked RyR1 particles accompanied by DHPR-like densities and then performed STA with C4 symmetry (**Fig. S2b**), resulting in a final cryo-EM map of the RyR1-DHPR supercomplex with a resolution of 33 Å (**Fig. S3d**), which features RyR1 on the TCM and DHPRs on the opposing TTM (**Fig. 5d and Movie S8**). The alignment procedure using C1 symmetry yielded a similar map with four extruding densities on TTM, showing that the DHPRs were arranged in a nearly C4 symmetrical pattern corresponding to the RyR1 tetramer (**Fig. S9a**). The extracellular domains (ECDs) of DHPRs within the T-tubular lumen resemble the tetrad that was observed on TTMs in previous freeze-fracture studies (*29*) and can be fitted well with the model of the α2δ subunit of reported DHPR structures (PDB entry 5GJW) (*63*) in terms of both shape and size (**Figs. S9b-c**).

To further investigate the direct interaction between RyR1 and DHPR, we fitted the above full-length mouse RyR1 model and the previously reported DHPR structure (PDB entry 5GJW) into the map and derived the structural model of the RyR1-DHPR supercomplex (**Fig. 5e and Movie S8**). In this model, the main body of each DHPR molecule, consisting of α1, α2δ and γ subunits, lies on top of the handle region of RyR1 instead of the RyR1 tetramer corner (**Fig. 5f**). Notably, the densities of RyR1 and DHPR exhibit two direct connections that are observed in both raw tomograms (**Fig. 5b**) and the averaged map (**Fig. 5g**). The two connection densities, denoted as Connections 1 and 2, are found near the P1 and P2 subdomains of RyR1, respectively (**Figs. 5h-i**).

### Arrangement of the RyR1-DHPR supercomplex in the native triad junction

By plotting the aligned RyR1-DHPR map back to the corresponding tomograms, we found that tetrads do not appear opposite every RyR1. Instead, they tend to skip one of two adjacent RyR1s in the same row and bind to the other RyR1 (**Figs. 6a and Movie S8**). This result is consistent with our statistics of the distances between two neighbouring RyR1-DHPR supercomplexes (**Figs. 6b**). Two peaks at ∼44 nm and ∼62 nm were found in the distribution histogram. The 44 nm peak coincides with the distance between two adjacent RyR1-DHPR complexes from different rows, and the peak of 62 nm matches the distance between two adjacent complexes in the same row (**Fig. 6c**). This unique RyR1-DHPR arrangement provides a good explanation for the previous observation in freeze-fracture experiments where the dispositions of tetrads in two adjacent rows were found in an alternating way (*29*).

**Figure 6.**
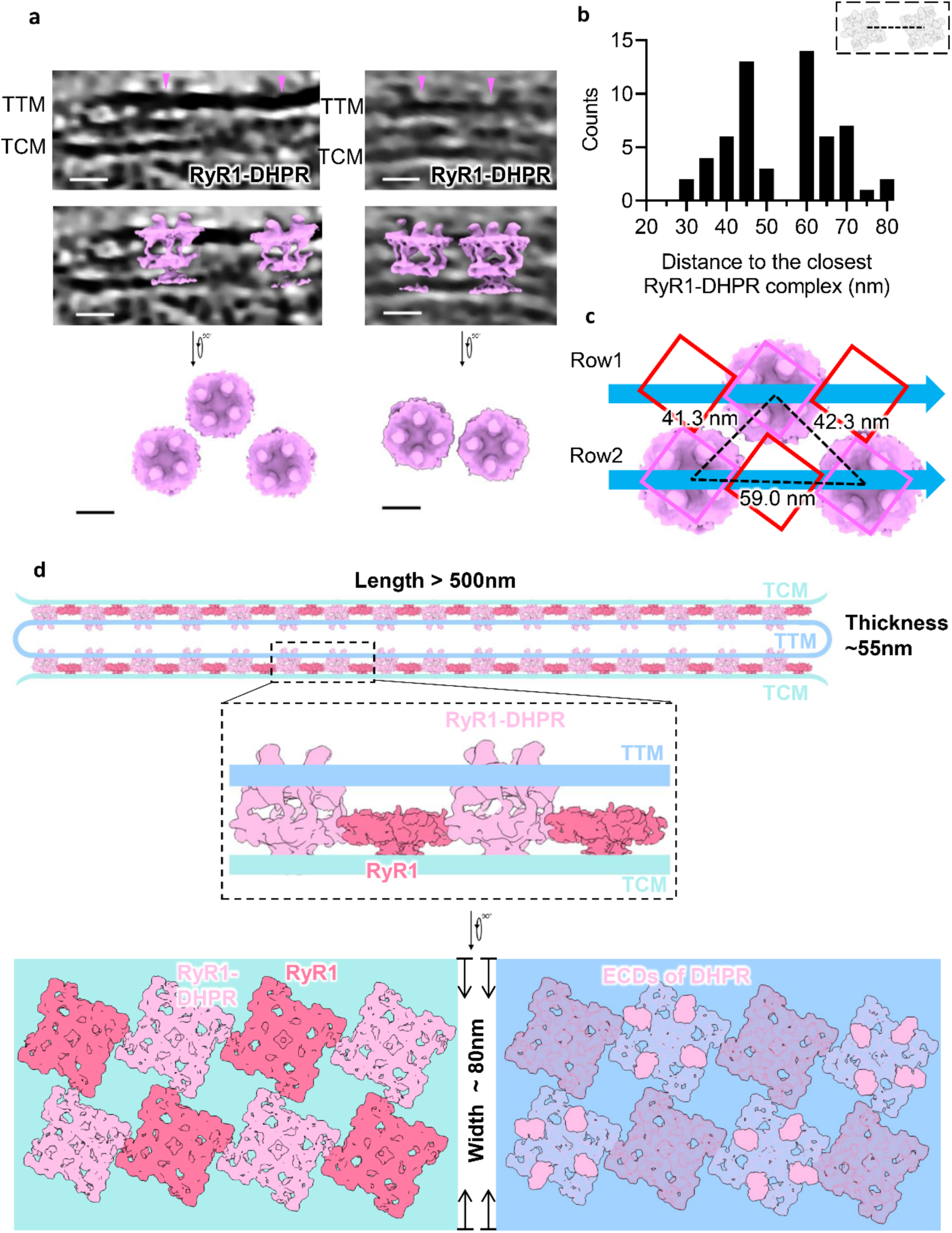
The arrangement of the RyR1-DHPR supercomplex in the native triad junction. (**a**) Representative tomogram slice of the adjacent RyR1-DHPR supercomplex in one triad junction. The STA averaged maps of RyR1-DHPR supercomplexes are plotted back into the tomograms and shown as the side and top views, respectively. Scale bar, 20 nm. (**b**) The distribution histogram of distances between adjacent RyR1-DHPR supercomplexes in the tomograms. (**c**) The arrangement of RyR1-DHPR supercomplexes in the triad junction with their nearest distances measured. The locations of RyR1 tetramers without DHPR are indicated as red squares. (**d**) The architecture of the native triad junction is shown as a diagram with TCM and TTM coloured light cyan and light blue, respectively. RyR1s without DHPR bound are shown in pink, and the RyR1-DHPR supercomplex is shown in pale magenta. The coordinates of the T-tubule are indicated. The top view of TCM from the cytoplasmic side and the top view of TTM from the lumen side of the T-tubule are shown below with the TTM set to partial transparency to show the arrangement of RyR1s.

By combining the above observations of arrangements of RyR1s and RyR1-DHPR supercomplexes as well as the statistical measurement of triad junction morphology, we were able to build an architectural model of the skeleton muscle triad junction (**Fig. 6d**). The entire triad junction is characterized by a long and narrow T-tubule sandwiched by two TCMs. The average length of the triad junction can be over 500 nm with a width of ∼80 nm and a thickness of ∼ 55 nm. The width of the triad junction can accommodate only two rows of the RyR1 array embedded in the TCM. The distance between TCM and TTM is similar to the interstitial space of the triad junction, which allows the interaction between the cytoplasmic region of RyR1 and that of DHPR. At the TCM, RyR1s interact with each other in a right-hand corner-to-corner way to form a two-row array. At the TTM, every four DHPRs form a tetrad that interacts with the handle regions of the RyR1 tetramer at the interstitial space, forming the RyR1-DHPR supercomplex. The tetrads of DHPR also form an ordered arrangement at the TTM; however, unlike the array of RyR1s, these tetrads are not proximal to each other because one tetrad appears every two RyR1 tetramers. Therefore, the RyR1-DHPR supercomplexes and RyR1s without bound DHPR are arranged in an alternating checkboard-like pattern within the triad junction.

## Discussions

In the present study, to elucidate the molecular mechanism of ECC of skeletal muscle, we resolved the native architecture of the mouse skeletal muscle triad junction at nanometre resolution by utilizing *in situ* cryo-ET techniques. With subtomogram analysis, the *in situ* structure of RyR1 in complex with FKBP12 and apo-CaM was solved to 16.7 Å and found to form a two-row array with a right-hand corner-to-corner interaction pattern on the TCM. Furthermore, DHPRs were found on the TTM, and every four DHPRs formed a tetrad that interacted with the RyR1 tetramer in an alternating way. From the *in situ* structure of the RyR1-DHPR supercomplex, two direct contact sites between RyR1 and DHPR were observed, with one located at the P1 domain of RyR1 and another located at the P2 domain. Overall, the RyR1-DHPR supercomplexes and RyR1s without DHPR bound are arranged in an alternating checkboard-like pattern within the triad junction. To understand the biological importance of the right-hand corner-to-corner arrangement of RyR1s, we performed molecular dynamics simulations and found that this unique arrangement favours strong interactions among RyR1s in the open state, thus enabling synergic calcium release within the array of RyR1s.

ECC of muscle has been proposed for many years, and a vast amount of research has been performed in decades to explore its molecular mechanism. At the triad junction, the coupling procedure includes two steps: the first step is the coupling between the activation of DHPR and that of RyR, and the second step is the coupling of the synergic calcium release of neighbouring RyRs.

For the first step of coupling where DHPR is involved, two different mechanisms were proposed. One proposed mechanism was chemical coupling between DHPR and RyR2 in cardiac muscle, in which the calcium released by the voltage-gated DHPR triggers the subsequent activation of RyR2 (*66*). The second proposed mechanism was physical coupling between DHPR and RyR1 in skeletal muscle (*29*), but this suggestion lacked direct evidence. Our results included direct observation of the physical interaction between DHPR and RyR1 with two connection sites, Connections 1 and 2. The α1 subunit of DHPR plays a crucial role in the ECC process. It not only functions as a voltage sensor but also plays a potential role in delivering signals to RyR1 directly. We predicted the complete structural model of the DHPR α1 subunit using AlphaFold2 (*67*) and found that this subunit could potentially occupy the density of Connection 2 by its Ⅱ-Ⅲ loop between the II and III repeats and its C-terminal domain (CTD) and then make direct contact with RyR1 (**Fig. S9d**). This is consistent with previous studies showing that the Ⅱ-Ⅲ loop might interact directly with RyR1 (*38, 68*) and is critical for ECC (*31*). Fluorescence resonance energy transfer (FRET) experiments also showed that the Ⅱ-Ⅲ loop undergoes conformational changes during the action potential, possibly transmitting signals to RyR1 (*40*).

STAC3 (SH3 and Cysteine Rich Domain 3) was predicted to be a crucial component for the interaction between RyR1 and DHPR (*69, 70*). Its PKC (protein kinase C-like) domain has been shown to interact with the CTD and II-III loop of the α1 subunit (*71, 72*). Additionally, junctophilin2 (JP2) was reported as a minimum requirement for the ECC process, and its N-terminus was found to bind to the last 12 residues of the α1 subunit (*73, 74*). Therefore, it is highly likely that STAC3 and JP2 are also included in the formation of the RyR1-DHPR supercomplex and located at the sites of Connection 2.

Previous studies tried to identify the significant region of RyR1 for the ECC process by constructing the RyR1-RyR2 or RyR1-RyR3 chimeric proteins, and one of them found that residues 1272-1455 of RyR1, which belong to the SPRY3 domain, are crucial for the formation of the DHPR tetrad (*36*), suggesting its role lies in forming direct interactions with DHPR. Our study found that such a region is located near the density of Connection 1, which is consistent with the previous study. In addition, another study determined that residues 1635-3720 of RyR1 form the region required for ECC restoration (*34*), which is situated near the density of Connection 2 in our study. Therefore, both previous functional studies and our structural study suggest that RyR1 and DHPR engage in rich interaction modes and possess multiple binding sites to ensure efficient direct signal transduction during the ECC process, which will be further validated by a higher-resolution structural study of the RyR1-DHPR complex.

The second step of coupling was proposed previously (*75, 76*) based on the observation of an array localization pattern of RyR1/2 channels in cardiac cells (*25*) and muscle tissues (*59*) and the observation of a spontaneous local increase in the concentration of intracellular calcium, commonly known as “Ca^2+^ sparks” (*62*). Such coupling is important to facilitate the propagation of the gating signal of RyR1/2s and thereby trigger synchronized muscle contraction. In cardiac cells, the gating kinetics of RyR2 are believed to be related to its array-based interaction, as RyR2s tend to form large clusters in cardiac cells (*65*). This interaction within RyR2 arrays not only transitions the gating of RyR2 from a thermodynamically reversible mode to an irreversible mode but also decreases the opening time of RyR2 channels (*65*). In the triad junction of skeletal muscle, RyR1 tetramers are also well organized into two parallel rows with a right-hand corner-to-corner interaction mode, suggesting a potential array-based coupling mechanism consistent with RyR2s in cardiac cells. Our study solved such array-based interactions at nanometre resolution, which enabled us to build a reliable structural model of the RyR1 array and their interaction interfaces, providing a molecular explanation of previous disease-related mutations (*61*), which could potentially abolish the coupled gating among RyR1s, thereby attenuating the effect of “Ca^2+^ sparks“. In addition, our MD simulations allowed us to observe the synergistic conformational changes of neighbouring RyR1s in the open state, providing an in-depth understanding of the mechanism of array-based coupling by RyR1s.

The checkboard-like arrangement of RyR1-DHPR supercomplexes and RyR1s without DHPR bound is a unique architectural feature of the triad junction of skeleton muscle, and this result deepens our understanding of the ECC process. We speculate that there might be some structural hindrances from adjacent DHPR tetrads or allosteric unfavourable conformational changes of neighbouring RyR1 tetramers to prevent the formation of the RyR1-DHPR supercomplex for every RyR1 tetramer. Moreover, it is not necessary to form the RyR1-DHPR supercomplex for every RyR1 tetramer because the corner-to-corner interaction between adjacent RyR1 tetramers ensures synergistic gating once one neighbouring RyR1 tetramer is activated. In addition, the coupling efficiency by the physical corner-to-corner interaction would decrease along the propagation distance because the gating signal could not be amplified by the physical coupling in terms of the energy view. Therefore, the arrangement of DHPR tetrads for every two RyR1 tetramers is important to facilitate a fast and efficient ECC process and achieve spontaneous activation of RyR1s in the triad junction to ensure synergistic contraction of the entire fibril.

In conclusion, our study elucidated the inherent architecture of triad junctions in mammalian skeletal muscle and included the direct observation of the arrangement of RyR1 and the RyR1-DHPR supercomplex within the triad junction. This architecture facilitates a fast and efficient ECC process: a long and narrow triad junction network spreading over the long skeletal muscle fibres guarantees rapid signal transmission. Once a DHPR tetrad in the T-tubule is activated by voltage potential, it activates the RyR1 underneath instantly through physical coupling; the activated RyR1 then activates the neighbouring RyR1s instantly, also through physical coupling. In this way, a vast amount of calcium is released from the SR, and muscle contraction is induced. This intricate and ingenious construction of triad junctions ensures rapid signal transmission during the ECC process, thereby preserving the normal physiological functions of skeletal muscle.

## Methods

### Extraction and vitrification of mouse skeletal muscle fibres

The EDL muscles were separated from 2∼3-week-old mice and incubated in Dulbecco’s modified Eagle’s medium (DMEM) (high glucose, L-glutamine with 110 mg/L sodium pyruvate) containing 0.2% collagenase at 37 °C for digestion. During this time, the condition of muscles should be regularly checked to avoid overdigestion. After ∼30 minutes, when the muscles began to loosen, they were transferred to a new dish containing DMEM without collagenase to stop the digestion. Then, the pipette was used to gently flush the muscles with medium until the muscle was naturally released and the live muscle fibres were spread in the medium. During this process, the flushing time when the sample was exposed to room temperature was less than 10 minutes. Then, the dish was transferred into an incubator for incubation at 37 °C and 5% CO_2_ for at least 5 minutes to keep the muscle fibres alive before another round of flushing.

Before plunge freezing, the isolated single muscle fibre was incubated in medium containing 10% glycerol for 3-5 minutes to avoid incomplete vitrification. Then, 3 μl of medium containing one isolated muscle fibre was placed on the glow-discharged grid (Quantifoil R2/1 Au 200 mesh). After incubation for 60 seconds at 37 °C, the grid was blotted from the opposite side for 10 seconds using EMGP2 (Leica) and then plunge frozen into liquid ethane.

### Cryo-lamella preparation using cryo-FIB

The vitrified grid containing the isolated muscle fibre was clipped into a special AutoGrid (Thermo Fisher Scientific) designed for cryo-FIB milling. After being loaded into the transfer shuttle (*77*), the grid was transferred into a cryo-FIB/SEM dual-beam microscope Aquilos2 (Thermo Fisher Scientific). Before milling, the grid was sputter coated with platinum for 10 seconds to reduce the charging effects of the sample. Then, it was injected with organometallic platinum for 12 seconds to prevent damage by gallium ion beam milling. The grid was pretilted 8 degrees to expose the muscle fibres on the grid. The sample was milled to ∼300 nm following the regular steps. Then, the cryo-lamella was polished further using a 10 nA current and examined using cryo-SEM images to estimate the thickness by charging propensity. The process was continued until the thickness reached ∼120 nm.

### Tilt series acquisition

The grids were inserted into the autogrid cassette at an angle that was 90 degrees rotated relative to the angle when inserted into the cryo-FIB/SEM microscope to ensure that the lamellae milling direction was perpendicular to the tilt axis of the electron microscope stage so that the nearby sample that was not milled would not block the lamellae during rotation. Afterwards, the cassette was loaded into a transmission electron microscope Titan Krios G3 (Thermo Fisher Scientific) equipped with a zero-loss energy filter and a K2 Summit direct electron detector (Gatan). Serial EM software (*78*) was used to acquire images. Target lamellae containing areas of interest were mapped at a low magnification of 6,300x to check and select the collection sites. Tilt series were acquired at 64000x magnification (pixel size 2.22 Å) with a dose-symmetric tilt scheme using the beam-shift method (*79*). The total dose was set to 140 to 160 e^-^/Å^2,^ and the tilt range was approximately -40° to +40° relative to the lamella pretilt angle (8°) with an increment of 2°. The actual range may be adjusted according to the condition of each lamella. The defocus was set to range between 4 μm and 5 μm.

### Image processing

Raw images of the tilt series were sent to Warp (*80*) for motion correction and CTF estimation. After manually filtering out the bad tilt images, the preprocessed tilt series were aligned using the patch-tracking method included in the IMOD software package (*81*). Then, the alignment parameters were sent back to Warp, and the tomograms were reconstructed. To visualize the densities of interest more clearly, we performed nonlinear anisotropic diffusion (NAD) filtering on the tomograms (*82*). Each RyR1 particle centre was manually picked in Dynamo (*83*), and its initial orientation was determined based on the vector between the manually selected RyR1 transmembrane region and cytoplasmic region.

A total of 4379 particles were picked from 264 tomograms. Then, relion3.1 (*84*) was utilized for the STA assay. First, the subtomogram particles in a box size of 48^3^ voxels were extracted from 8× binned tomograms in Warp. The direct reconstruction map using predetermined orientations was used as the initial reference for alignment. C4 symmetry was always applied, and a mask surrounding the cytoplasmic region of RyR1 was used to help align. After global refinement, a 3D classification was applied, and a better class containing 51% of the particles (2241) was selected for subsequent refinement. After re-extracting from 2× binned tomograms with a box size of 128^3^ voxels, a local refinement was performed, and a 3D classification followed. Then, 61% of particles (1365) in a better class were used for final local refinement, and the resolution was estimated to be 16.8 Å based on the ‘‘gold-standard’’ Fourier shell correlation (FSC) with a 0.143 criterion after postprocessing.

For the RyR1-DHPR supercomplex, 1421 particles were manually picked from 220 tomograms. First, a box size of 48^3^ voxels was used to extract the particles from 8× binned tomograms in Warp. The low-pass filtered RyR1 density map was used as the initial reference for alignment. Both C4 and C1 symmetry were applied during the process, confirming the four extruding densities for DHPR. A 3D classification without a mask was performed after global refinement, and a class containing 18% particles (251) was selected for local refinement. This density map contained four extruding densities assumed to be DHPRs. Then, all 1421 particles were extracted using a box size of 96^3^ voxels from 4× binned tomograms. After local refinement, a mask surrounding the RyR1 and DHPR α2δ subunit densities was used in the 3D classification procedure, and a class containing 11% particles (154) was selected from 8 classes. The density map containing one RyR1 and four DHPRs was finally obtained after local refinement and postprocessing.

### Model building

The preliminary model of mouse RyR1 was forecasted using the SWISS-MODEL homology modelling method (*85*). Alphafold2 was used to predict the partial structure of RyR1 because the sequence exceeded the maximum allowed by homology modelling. Subsequently, the complete structure of RyR1 was finalized using Coot software (*86*). The models of RyR1, FKBP12 and CaM are fitted in the reconstruction map using chimaera (*87*).

Then, we performed stepwise MDFF simulations to refine the complex model according to the density map. A time step of 1 fs was used throughout the simulation. Langevin dynamics were adopted at a temperature of 310 K. The equilibration step for energy minimization was performed on the initial model for 1000 steps before the refinement run. The refinement runs were performed for 3,000 ps, corresponding to 3,000,000 simulation steps, and the gridForceScale values were gradually increased from 0.3 to 0.7 during the refinement. All simulations were performed using CHARMM36m force fields (*88*). Electrostatic calculations were treated with particle mesh Ewald (PME). A cut-off of 12 Å was chosen for short-range van der Waals interactions. NAMD (*89*) was used as the MD engine throughout all simulations.

### Molecular dynamics simulation

The open and closed RyR1 structures were constructed using homology modelling in Maestro (Schrödinger, LLC) with the sequence information and major templates displayed in **Table S2**; missing segments were built using the trRosetta server (*90*). The membrane builder in CHARMM-GUI (*91*) was employed to build the protein-membrane complex system. To mimic the endoplasmic reticulum (ER) membrane, the membrane for RyR1 consisted of 54% phosphocholine (POPC), 18% phosphatidylethanolamine (POPE), 7% cholesterol, 5% phosphatidylserine (POPS), 8% phosphatidylinositol (POPI), and 5% N-palmitoylsphingomyelin (PSM) in the molar ratio (*92*). The protein-membrane complex was solvated in an aqueous phase containing TIP3P water molecules, K^+^ or Cl^-^ as counter ions, and 0.05 M CaCl_2_ in a periodic box. A system with one RyR1 was first constructed and equilibrated, then further built into a two-RyR1 system using VMD (*93*). All-atom RyR1 systems contain up to six million atoms, which are computationally expensive to simulate on a long time scale. All-atom (AA) simulations were applied to explore the interactions and motions of two-RyR1 assembled by intermolecular helical-helical domain interactions, while CG simulations were used to explore the assembly of the four-RyR1 system.

For CG simulations, we used the Martini Bilayer Maker in CHARMM-GUI (*91*) to construct the protein-membrane complex systems, each of which contains CG models of ion channels, a CG bilayer with similar lipid components as in the AA systems, the nonpolarizable water models, Na^+^ or Cl^-^ as counter ions, and 0.05 M NaCl. A system with one RyR1 was first constructed and equilibrated, which was further built into a four-RyR1 system using VMD (*93*). A summary of the simulation systems is provided in **Table S3**.

All-atom simulations were performed with the CHARMM36m-cmap force field (*94*). Each system was subjected to energy minimization, 2 ns multistep equilibration under isothermal-isovolumetric (NVT) and isothermal-isobaric ensemble with semi-isotropic pressure coupling (NPγT) with the force constant of dihedral restraint on proteins and lipids gradually reduced from 250 kcal/mol to 0 kcal/mol in six steps. Then, the system continued to relax in MD simulations under the NPγT ensemble (310 Kelvin, 1 bar, Langevin dynamics thermostat and semi-isotropic Monte Carlo barostat). A time step of 2 fs was used for the initial 7 ns relaxation and then increased to 3 fs under hydrogen mass repartitioning. All lengths of bonds to hydrogen atoms in protein or lipid molecules were constrained with SHAKE. The PME technique (*95*) was used to calculate long-range electrostatic interactions. The van der Waals and short-range electrostatics were cut off at 12.0 Å with a switch at 10.0 Å. The CG simulations were carried out with Martini 2.2 with an elastic network (elnedyn22) force field using the Gromacs 2019 program (*96*) with a time step of 20 fs under the NPγT ensemble (310 Kelvin, 1 bar, v-rescale thermostat and Parrinello-Rahman barostat).

Trajectory analyses such as distances, short-distance contacts, hydrogen bonds, PCA, and dynamic cross-correlation map (DCCM) were performed with “cpptraj” (*97*) and then further processed and plotted using matplotlib (*98*). Structures were shown by PyMOL (Schrödinger, LLC), and movies were generated in VMD (*93*). Solvent-accessible pore radii along a channel were calculated by HOLE (*99*) and visualized and rendered in VMD. The intermolecular helical-helical domain with a minimum distance less than 8 Å at the end of 3 μs CG simulations is considered to form a right-hand contact. We counted the contact numbers from ten copies. The numbers of all contacts (< 6 Å), H-bond pairs, and hydrophobic contacts (< 6 Å) were counted from the last 110 ns of all three all-atom simulations for the open and closed RyR1-RyR1 systems. For both the PCA and DCCM analyses, we used the trajectories from the last 110 ns of all three copies. We first aligned the Cα of the interacting RyR1-RyR1 structure on the first frame by the sequences that are largely nonloop (i.e., 12-1297, 1432-1871, 1924-2042, 2092-2826, 2852-4255, 4541-4576, 4639-5035 of each RyR1 monomer) to generate an averaged coordinate, on which we aligned again and then used it for the PCA and DCCM analyses. The cross-correlation matrix elements, *C*_ij_, are defined as (*100*):

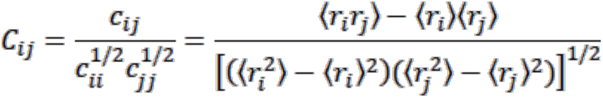

where *i* and *j* represent two atoms with the position vectors in the (fitted) structure at time *t* as *r*_i_(*t*) and *r*_j_(*t*), respectively; angle brackets denote time averages; and *c*_ij_ represents the corresponding covariance matrix element.

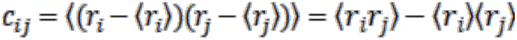

The values of the cross-correlation coefficients range from -1 (completely anticorrelated motions) to 0.0 (no correlation) to 1.0 (completely correlated motions). The DCCM of the two interacting RyR1s at the contact interface or most distant RyR1s were studied. We calculated the first principal component of residues 2494-3611 (which contains the P2 domain and HD2 domain) from dimerized RyR1 in both open and closed states.

## Data availability

The raw tilt series used in this study has been deposited in EMPIAR (the Electron Microscopy Public Image Archive) China (http://www.emdb-china.org.cn) under accession code EMPIARC-XXXXX. The sub-tomogram averaged cryo-EM maps of RyR1 tetramer, dimeric RyR1 tetramers, and RyR1-DHPR supercomplex in C1 and C4 symmetries have been deposited in the Electron Microscopy Database (EMDB) with the accession codes EMD-37089, EMD-37092, EMD-37093 and EMD-37094, respectively. Other data that support the findings of this study are available from the corresponding author upon request.

## Acknowledgments

We thank Ping Shan, Lianwan Chen, and Tongming Zhang (F.S. laboratory) for their assistance in the laboratory management. We thank the Center for Biological Imaging, Institute of Biophysics, Chinese Academy of Science for the cryo-EM work, and we are grateful to Dr. Shuoguo Li, Lulu Qin, and Dr. Jianguo Zhang for their help with the cryo-EM sample preparation and data collection.

This work was equally supported by grants from the National Natural Science Foundation of China for Distinguished Young Scholars (31925026 to FS), Chinese National Key Research and Development Program (2021YFA1301500 to FS and 2019YFA0709400 to GL), and the Strategic Priority Research Program of Chinese Academy of Sciences (XDB 37040102 to FS and XDB37040401 to GL). This work was also supported by grants from Chinese National Key Research and Development Program (2019YFA0904101 to YZ and 2017YFA0504702 to CCY) and National Natural Science Foundation of China (32071187 to YZ, 21933010 to GL, 22207108 to GL, 31830020 to FS, and 31770785 to YCC).

## Author contributions

F.S. conceived and supervised the study; F.S., and Y.Z. designed and coordinated the research; J.X. performed all the cryo-EM experiments; C.L. performed all the molecular simulation experiments; J.X., C.L., G.L. and Y.Z. analyzed the data; J.X., C.L. G.L. and Y.Z. wrote the manuscript; F.S., Y.Z. and C.C.Y. revised the manuscript.

## References

1. A. Sandow, Excitation-contraction coupling in muscular response. The Yale journal of biology and medicine 25, 176–201 (1952).

2. H. E. Huxley, Evidence for Continuity Between the Central Elements of the Triads and Extracellular Space in Frog Sartorius Muscle. Nature 202, 1067–1071 (1964).

3. H. González-Serratos, Inward spread of activation in vertebrate muscle fibres. J Physiol 212, 777–799 (1971).

4. C. Franzini-Armstrong, K. R. Porter, SARCOLEMMAL INVAGINATIONS CONSTITUTING THE T SYSTEM IN FISH MUSCLE FIBERS. The Journal of cell biology 22, 675–696 (1964).

5. M. F. Schneider, W. K. Chandler, Voltage Dependent Charge Movement in Skeletal Muscle: a Possible Step in Excitation–Contraction Coupling. Nature 242, 244–246 (1973).

6. E. Ríos, G. Brum, Involvement of dihydropyridine receptors in excitation–contraction coupling in skeletal muscle. Nature 325, 717–720 (1987).

7. B. A. Adams, T. Tanabe, A. Mikami, S. Numa, K. G. Beam, Intramembrane charge movement restored in dysgenic skeletal muscle by injection of dihydropyridine receptor cDNAs. Nature 346, 569–572 (1990).

8. O. Delbono, E. Stefani, Calcium transients in single mammalian skeletal muscle fibres. J Physiol 463, 689–707 (1993).

9. H. Takeshima et al., Primary structure and expression from complementary DNA of skeletal muscle ryanodine receptor. Nature 339, 439–445 (1989).

10. M. B. Bhat, J. Zhao, H. Takeshima, J. Ma, Functional calcium release channel formed by the carboxyl-terminal portion of ryanodine receptor. Biophys J 73, 1329–1336 (1997).

11. A. Saito, M. Inui, M. Radermacher, J. Frank, S. Fleischer, Ultrastructure of the calcium release channel of sarcoplasmic reticulum. The Journal of cell biology 107, 211–219 (1988).

12. J. Nakai et al., Primary structure and functional expression from cDNA of the cardiac ryanodine receptor/calcium release channel. FEBS letters 271, 169–177 (1990).

13. K. Otsu et al., Molecular cloning of cDNA encoding the Ca2+ release channel (ryanodine receptor) of rabbit cardiac muscle sarcoplasmic reticulum. J Biol Chem 265, 13472–13483 (1990).

14. Y. Hakamata, J. Nakai, H. Takeshima, K. Imoto, Primary structure and distribution of a novel ryanodine receptor/calcium release channel from rabbit brain. FEBS letters 312, 229–235 (1992).

15. C. Franzini-Armstrong, STUDIES OF THE TRIAD : I. Structure of the Junction in Frog Twitch Fibers. The Journal of cell biology 47, 488–499 (1970).

16. F. A. Lai, K. Anderson, E. r. Rousseau, Q.-Y. Liu, G. Meissner, Evidence for a Ca2+ channel within the ryanodine receptor complex from cardiac sarcoplasmic reticulum. Biochemical and biophysical research communications 151 **1**, 441–449 (1988).

17. T. Wagenknecht et al., Three-dimensional architecture of the calcium channel/foot structure of sarcoplasmic reticulum. Nature 338, 167–170 (1989).

18. R. G. Efremov, A. Leitner, R. Aebersold, S. Raunser, Architecture and conformational switch mechanism of the ryanodine receptor. Nature 517, 39–43 (2015).

19. Z. Yan et al., Structure of the rabbit ryanodine receptor RyR1 at near-atomic resolution. Nature 517, 50–55 (2015).

20. R. Zalk et al., Structure of a mammalian ryanodine receptor. Nature 517, 44–49 (2015).

21. A. des Georges et al., Structural Basis for Gating and Activation of RyR1. Cell 167, 145–157 e117 (2016).

22. R. Wei et al., Structural insights into Ca(2+)-activated long-range allosteric channel gating of RyR1. Cell Res 26, 977–994 (2016).

23. C. Franzini-Armstrong, J. W. Kish, Alternate disposition of tetrads in peripheral couplings of skeletal muscle. J Muscle Res Cell Motil 16, 319–324 (1995).

24. C. Franzini-Armstrong, The sarcoplasmic reticulum and the control of muscle contraction. Faseb j 13 **Suppl 2**, S266–270 (1999).

25. C. Franzini-Armstrong, F. Protasi, V. Ramesh, Shape, size, and distribution of Ca(2+) release units and couplons in skeletal and cardiac muscles. Biophys J 77, 1528–1539 (1999).

26. K. E. Loesser, L. Castellani, C. Franzini-Armstrong, Dispositions of junctional feet in muscles of invertebrates. J Muscle Res Cell Motil 13, 161–173 (1992).

27. D. G. Ferguson, H. W. Schwartz, C. Franzini-Armstrong, Subunit structure of junctional feet in triads of skeletal muscle: a freeze-drying, rotary-shadowing study. The Journal of cell biology 99, 1735–1742 (1984).

28. C. C. Yin, L. M. Blayney, F. A. Lai, Physical coupling between ryanodine receptor-calcium release channels. J Mol Biol 349, 538–546 (2005).

29. B. A. Block, T. Imagawa, K. P. Campbell, C. Franzini-armstrong, Structural evidence for direct interaction between the molecular components of the transverse tubule/sarcoplasmic reticulum junction in skeletal muscle. The Journal of Cell Biology 107, 2587 - 2600 (1988).

30. C. Paolini, J. D. Fessenden, I. N. Pessah, C. Franzini-Armstrong, Evidence for conformational coupling between two calcium channels. Proc Natl Acad Sci U S A 101, 12748–12752 (2004).

31. T. Tanabe, K. G. Beam, B. A. Adams, T. Niidome, S. Numa, Regions of the skeletal muscle dihydropyridine receptor critical for excitation-contraction coupling. Nature 346, 567–569 (1990).

32. J. Nakai, N. Sekiguchi, T. A. Rando, P. D. Allen, K. G. Beam, Two regions of the ryanodine receptor involved in coupling with L-type Ca2+ channels. J Biol Chem 273, 13403–13406 (1998).

33. C. M. Wilkens, N. Kasielke, B. E. Flucher, K. G. Beam, M. Grabner, Excitation-contraction coupling is unaffected by drastic alteration of the sequence surrounding residues L720-L764 of the alpha 1S II-III loop. Proc Natl Acad Sci U S A 98, 5892–5897 (2001).

34. C. Proenza et al., Identification of a region of RyR1 that participates in allosteric coupling with the alpha(1S) (Ca(V)1.1) II-III loop. J Biol Chem 277, 6530–6535 (2002).

35. W. Cheng, X. Altafaj, M. Ronjat, R. Coronado, Interaction between the dihydropyridine receptor Ca2+ channel beta-subunit and ryanodine receptor type 1 strengthens excitation-contraction coupling. Proc Natl Acad Sci U S A 102, 19225–19230 (2005).

36. D. C. Sheridan et al., Bidirectional signaling between calcium channels of skeletal muscle requires multiple direct and indirect interactions. Proc Natl Acad Sci U S A 103, 19760–19765 (2006).

37. J. Schredelseker, A. Dayal, T. Schwerte, C. Franzini-Armstrong, M. Grabner, Proper restoration of excitation-contraction coupling in the dihydropyridine receptor beta1-null zebrafish relaxed is an exclusive function of the beta1a subunit. J Biol Chem 284, 1242–1251 (2009).

38. H. S. Tae et al., Molecular recognition of the disordered dihydropyridine receptor II-III loop by a conserved spry domain of the type 1 ryanodine receptor. Clin Exp Pharmacol Physiol 36, 346–349 (2009).

39. R. T. Rebbeck et al., The beta(1a) subunit of the skeletal DHPR binds to skeletal RyR1 and activates the channel via its 35-residue C-terminal tail. Biophys J 100, 922–930 (2011).

40. A. Polster, J. D. Ohrtman, K. G. Beam, S. Papadopoulos, Fluorescence resonance energy transfer (FRET) indicates that association with the type I ryanodine receptor (RyR1) causes reorientation of multiple cytoplasmic domains of the dihydropyridine receptor (DHPR) alpha(1S) subunit. J Biol Chem 287, 41560–41568 (2012).

41. A. Dayal, V. Bhat, C. Franzini-Armstrong, M. Grabner, Domain cooperativity in the β1a subunit is essential for dihydropyridine receptor voltage sensing in skeletal muscle. Proc Natl Acad Sci U S A 110, 7488–7493 (2013).

42. T. Wagenknecht, C.-e. Hsieh, M. Marko, Skeletal Muscle Triad Junction Ultrastructure by Focused-Ion-Beam Milling of Muscle and Cryo-Electron Tomography. European Journal of Translational Myology 25, (2015).

43. C. Renken et al., Structure of frozen-hydrated triad junctions: a case study in motif searching inside tomograms. J Struct Biol 165, 53–63 (2009).

44. W. Chen, M. Kudryashev, Structure of RyR1 in native membranes. EMBO Rep 21, e49891 (2020).

45. R. M. Sanchez, Y. Zhang, W. Chen, L. Dietrich, M. Kudryashev, Subnanometer-resolution structure determination in situ by hybrid subtomogram averaging - single particle cryo-EM. Nature Communications 11, 3709 (2020).

46. J. Zhang et al., VHUT-cryo-FIB, a method to fabricate frozen hydrated lamellae from tissue specimens for in situ cryo-electron tomography. J Struct Biol 213, 107763 (2021).

47. Z. Wang et al., The molecular basis for sarcomere organization in vertebrate skeletal muscle. Cell 184, 2135–2150 e2113 (2021).

48. R. Sousa, Use of glycerol, polyols and other protein structure stabilizing agents in protein crystallization. *Acta crystallographica. Section D*, Biological crystallography 51, 271–277 (1995).

49. J. H. Ye, J. Zhang, C. Xiao, J. Q. Kong, Patch-clamp studies in the CNS illustrate a simple new method for obtaining viable neurons in rat brain slices: glycerol replacement of NaCl protects CNS neurons. Journal of neuroscience methods 158, 251–259 (2006).

50. F. J. B. Bäuerlein et al., In Situ Architecture and Cellular Interactions of PolyQ Inclusions. Cell 171, 179–187.e110 (2017).

51. A.-M. B. Brillantes et al., Stabilization of calcium release channel (ryanodine receptor) function by FK506-binding protein. Cell 77, 513–523 (1994).

52. T. Wagenknecht et al., Locations of Calmodulin and FK506-binding Protein on the Three-dimensional Architecture of the Skeletal Muscle Ryanodine Receptor*. Journal of Biological Chemistry 272, 32463–32471 (1997).

53. A. Tripathy, L. Xu, G. Mann, G. Meissner, Calmodulin activation and inhibition of skeletal muscle Ca2+ release channel (ryanodine receptor). Biophys J 69, 106–119 (1995).

54. C. P. Moore et al., Apocalmodulin and Ca2+ Calmodulin Bind to the Same Region on the Skeletal Muscle Ca2+ Release Channel. Biochemistry 38, 8532–8537 (1999).

55. X. Huang, B. Fruen, D. T. Farrington, T. Wagenknecht, Z. Liu, Calmodulin-binding locations on the skeletal and cardiac ryanodine receptors. J Biol Chem 287, 30328–30335 (2012).

56. K. A. Woll, O. Haji-Ghassemi, F. Van Petegem, Pathological conformations of disease mutant Ryanodine Receptors revealed by cryo-EM. Nature Communications 12, 807 (2021).

57. Z. Melville et al., A drug and ATP binding site in type 1 ryanodine receptor. Structure (London, England : 1993) 30, 1025–1034.e1024 (2022).

58. D. Gong et al., Modulation of cardiac ryanodine receptor 2 by calmodulin. Nature 572, 347–351 (2019).

59. C. Franzini-Armstrong, G. Nunzi, Junctional feet and particles in the triads of a fast-twitch muscle fibre. J Muscle Res Cell Motil 4, 233–252 (1983).

60. L. M. Blayney et al., Ryanodine receptor oligomeric interaction: identification of a putative binding region. J Biol Chem 279, 14639–14648 (2004).

61. X. Chang et al., Correlation of Phenotype-Genotype and Protein Structure in RYR1-Related Myopathy. Front Neurol 13, 870285 (2022).

62. H. Cheng, W. J. Lederer, M. B. Cannell, Calcium sparks: elementary events underlying excitation-contraction coupling in heart muscle. Science 262, 740–744 (1993).

63. J. Wu et al., Structure of the voltage-gated calcium channel Ca(v)1.1 at 3.6 A resolution. Nature 537, 191–196 (2016).

64. R. Coronado et al., Functional equivalence of dihydropyridine receptor alpha1S and beta1a subunits in triggering excitation-contraction coupling in skeletal muscle. Biological research 37, 565–575 (2004).

65. S. Q. Wang, M. D. Stern, E. Rios, H. Cheng, The quantal nature of Ca2+ sparks and in situ operation of the ryanodine receptor array in cardiac cells. Proc Natl Acad Sci U S A 101, 3979–3984 (2004).

66. H. Cheng, S. Q. Wang, Calcium signaling between sarcolemmal calcium channels and ryanodine receptors in heart cells. Front Biosci 7, d1867–1878 (2002).

67. J. Jumper et al., Highly accurate protein structure prediction with AlphaFold. Nature 596, 583–589 (2021).

68. Y. Cui et al., A dihydropyridine receptor alpha1s loop region critical for skeletal muscle contraction is intrinsically unstructured and binds to a SPRY domain of the type 1 ryanodine receptor. Int J Biochem Cell Biol 41, 677–686 (2009).

69. E. J. Horstick et al., Stac3 is a component of the excitation-contraction coupling machinery and mutated in Native American myopathy. Nat Commun 4, 1952 (2013).

70. B. R. Nelson et al., Skeletal muscle-specific T-tubule protein STAC3 mediates voltage-induced Ca2+ release and contractility. Proc Natl Acad Sci U S A 110, 11881–11886 (2013).

71. M. Campiglio, B. E. Flucher, STAC3 stably interacts through its C1 domain with Ca(V)1.1 in skeletal muscle triads. Sci Rep 7, 41003 (2017).

72. S. M. Wong King Yuen, M. Campiglio, C. C. Tung, B. E. Flucher, F. Van Petegem, Structural insights into binding of STAC proteins to voltage-gated calcium channels. Proc Natl Acad Sci U S A 114, E9520–e9528 (2017).

73. S. Perni, M. Lavorato, K. G. Beam, De novo reconstitution reveals the proteins required for skeletal muscle voltage-induced Ca^2+^ release. Proceedings of the National Academy of Sciences 114, 13822–13827 (2017).

74. T. Nakada et al., Physical interaction of junctophilin and the Ca(V)1.1 C terminus is crucial for skeletal muscle contraction. Proc Natl Acad Sci U S A 115, 4507–4512 (2018).

75. S. O. Marx, K. Ondrias, A. R. Marks, Coupled gating between individual skeletal muscle Ca2+ release channels (ryanodine receptors). Science 281, 818–821 (1998).

76. X. F. Hu et al., Modulation of the oligomerization of isolated ryanodine receptors by their functional states. Biophys J 89, 1692–1699 (2005).

77. J. Zhang, G. Ji, X. Huang, W. Xu, F. Sun, An improved cryo-FIB method for fabrication of frozen hydrated lamella. J Struct Biol 194, 218–223 (2016).

78. D. N. Mastronarde, Automated electron microscope tomography using robust prediction of specimen movements. Journal of Structural Biology 152, 36–51 (2005).

79. C. Wu, X. Huang, J. Cheng, D. Zhu, X. Zhang, High-quality, high-throughput cryo-electron microscopy data collection via beam tilt and astigmatism-free beam-image shift. J Struct Biol 208, 107396 (2019).

80. D. Tegunov, P. Cramer, Real-time cryo-electron microscopy data preprocessing with Warp. Nature Methods 16, 1146–1152 (2019).

81. J. R. Kremer, D. N. Mastronarde, J. R. McIntosh, Computer Visualization of Three-Dimensional Image Data Using IMOD. Journal of Structural Biology 116, 71–76 (1996).

82. A. S. Frangakis, R. Hegerl, Noise Reduction in Electron Tomographic Reconstructions Using Nonlinear Anisotropic Diffusion. Journal of Structural Biology 135, 239–250 (2001).

83. D. Castaño-Díez, M. Kudryashev, M. Arheit, H. Stahlberg, Dynamo: A flexible, user-friendly development tool for subtomogram averaging of cryo-EM data in high-performance computing environments. Journal of Structural Biology 178, 139–151 (2012).

84. T. A. M. Bharat, S. H. W. Scheres, Resolving macromolecular structures from electron cryo-tomography data using subtomogram averaging in RELION. Nature Protocols 11, 2054–2065 (2016).

85. A. Waterhouse et al., SWISS-MODEL: homology modelling of protein structures and complexes. Nucleic acids research 46, W296–w303 (2018).

86. P. Emsley, B. Lohkamp, W. G. Scott, K. Cowtan, Features and development of Coot. Acta crystallographica. Section D, Biological crystallography 66, 486–501 (2010).

87. E. F. Pettersen et al., UCSF Chimera--a visualization system for exploratory research and analysis. Journal of computational chemistry 25, 1605–1612 (2004).

88. J. Huang et al., CHARMM36m: an improved force field for folded and intrinsically disordered proteins. Nat Methods 14, 71–73 (2017).

89. J. C. Phillips et al., Scalable molecular dynamics with NAMD. Journal of computational chemistry 26, 1781–1802 (2005).

90. Z. Du et al., The trRosetta server for fast and accurate protein structure prediction. Nature Protocols 16, 5634–5651 (2021).

91. S. Jo, T. Kim, V. G. Iyer, W. Im, CHARMM-GUI: a web-based graphical user interface for CHARMM. Journal of computational chemistry 29, 1859–1865 (2008).

92. B. Marzoog, T. Vlasova, Membrane lipids under norm and pathology. (2021).

93. W. Humphrey, A. Dalke, K. Schulten, VMD: visual molecular dynamics. J. Mol. Graph. Model. 14, 33–38 (1996).

94. R. B. Best et al., Optimization of the Additive CHARMM All-Atom Protein Force Field Targeting Improved Sampling of the Backbone ϕ, ψ and Side-Chain χ1 and χ2 Dihedral Angles. J. Chem. Theory Comput. 8, 3257–3273 (2012).

95. T. Darden, D. York, L. Pedersen, Particle Mesh Ewald - an N.Log(N) Method for Ewald Sums in Large Systems. J. Chem. Phys. 98, 10089–10092 (1993).

96. H. J. C. Berendsen, D. Vanderspoel, R. Vandrunen, Gromacs - a Message-Passing Parallel Molecular-Dynamics Implementation. Comput. Phys. Commun. 91, 43–56 (1995).

97. D. R. Roe, T. E. Cheatham, PTRAJ and CPPTRAJ: Software for Processing and Analysis of Molecular Dynamics Trajectory Data. J. Chem. Theory Comput. 9, 3084–3095 (2013).

98. J. D. Hunter, Matplotlib: A 2D graphics environment. Computing in Science & Engineering 9, 90–95 (2007).

99. O. S. Smart, J. G. Neduvelil, X. Wang, B. A. Wallace, M. S. P. Sansom, HOLE: A program for the analysis of the pore dimensions of ion channel structural models. Journal of Molecular Graphics 14, 354–360 (1996).

100. P. H. Hünenberger, A. E. Mark, W. F. van Gunsteren, Fluctuation and cross-correlation analysis of protein motions observed in nanosecond molecular dynamics simulations. J Mol Biol 252, 492–503 (1995).

101. I. Belevich, M. Joensuu, D. Kumar, H. Vihinen, E. Jokitalo, Microscopy Image Browser: A Platform for Segmentation and Analysis of Multidimensional Datasets. PLOS Biology 14, e1002340 (2016).

