## Supplementary Figures and Tables for "*In situ* structural insights into the excitation contraction coupling mechanism of skeletal muscle"

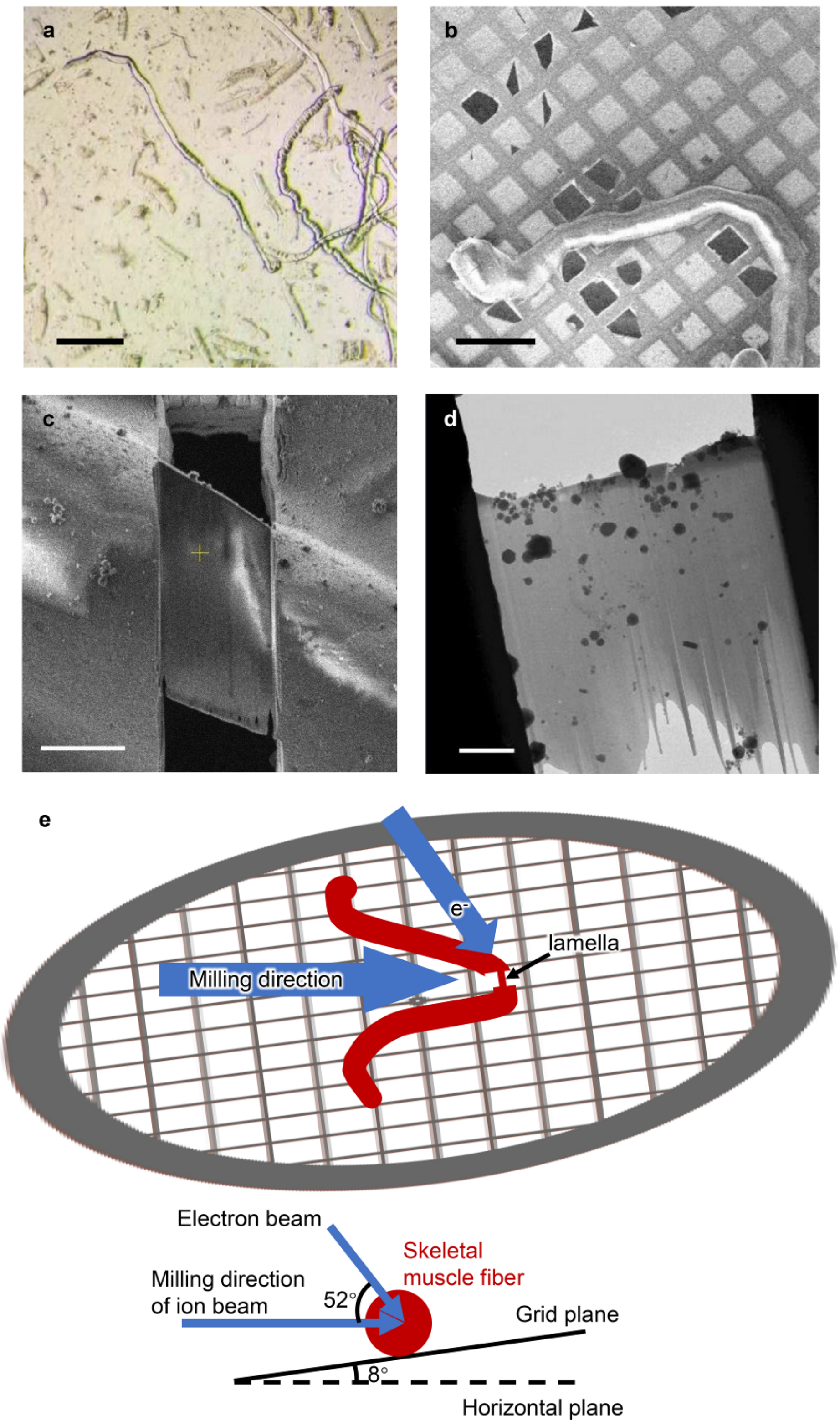

**Figure S1. Workflow of the sample preparation.** (**a**) Isolated mouse skeletal muscle fibers under light microscope. Scale bar, 300 μm. (**b**) Cryo-SEM image of the muscle fiber on the grids. Scale bar, 200 μm. (**c**) Cryo-SEM image of the cryo-lamella of muscle fiber after cryo-FIB milling. Scale bar, 10 μm. (**d**) Low magnification cryo-EM image of the cryo-lamella of muscle fiber. Scale bar, 2 μm. (**e**) Cartoon diagram shows the relative spatial relation between the ion beam and the skeletal muscle fiber (red).

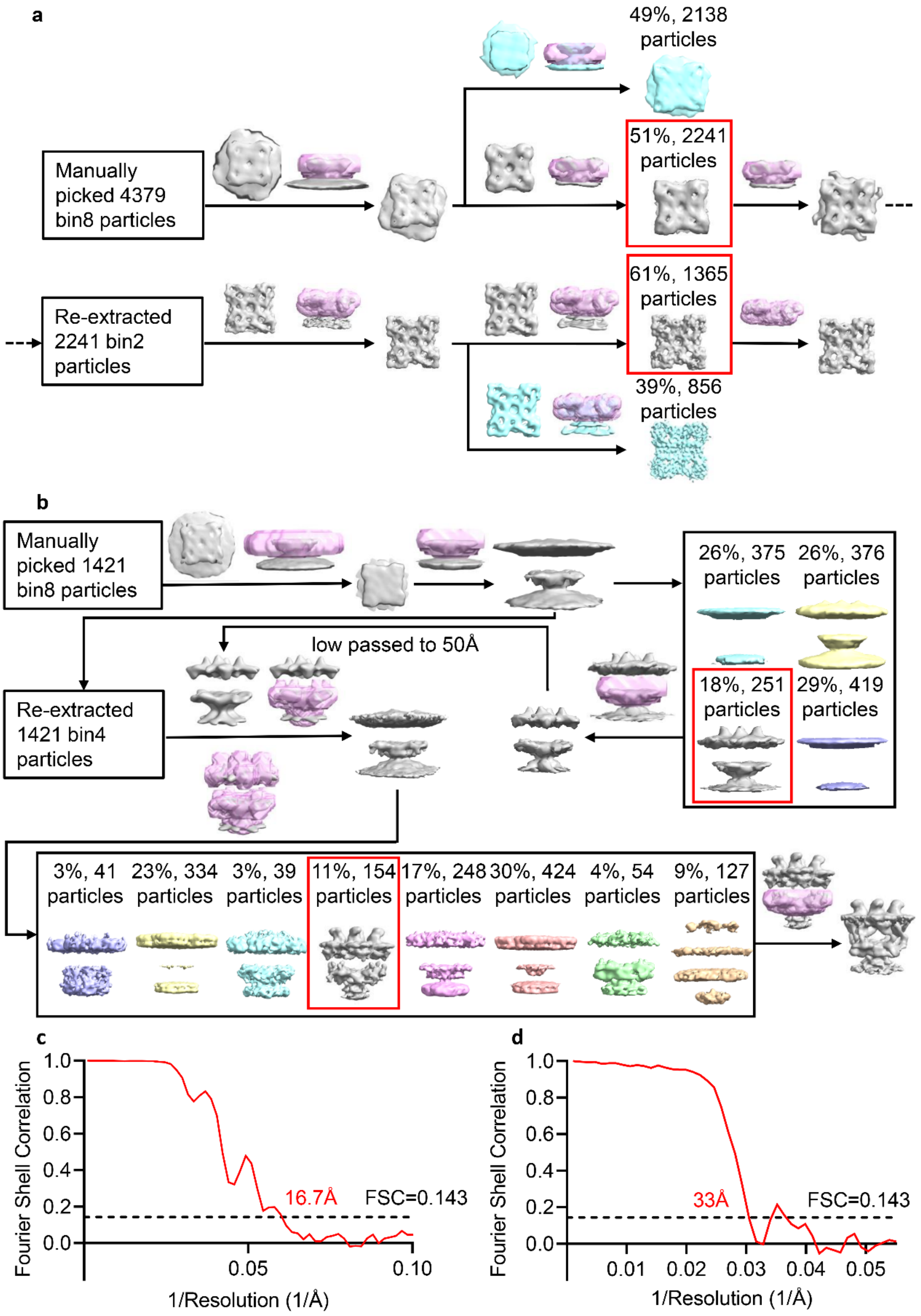


**Figure S2. Workflow of STA process for RyR1 (a) and RyR1-DHPR supercomplex (b).** The masks used during data process are indicated in pink. (**c**) The gold standard FSC curve of two half STA averaged maps of *in situ* RyR1. The estimated resolution of the reconstruction is 16.7 Å according to the 0.143 criterion. (**d**) The gold standard

FSC curve of two half STA averaged maps of *in situ* RyR1-DHPR supercomplex. The estimated resolution of the reconstruction is 33 Å according to the 0.143 criterion.

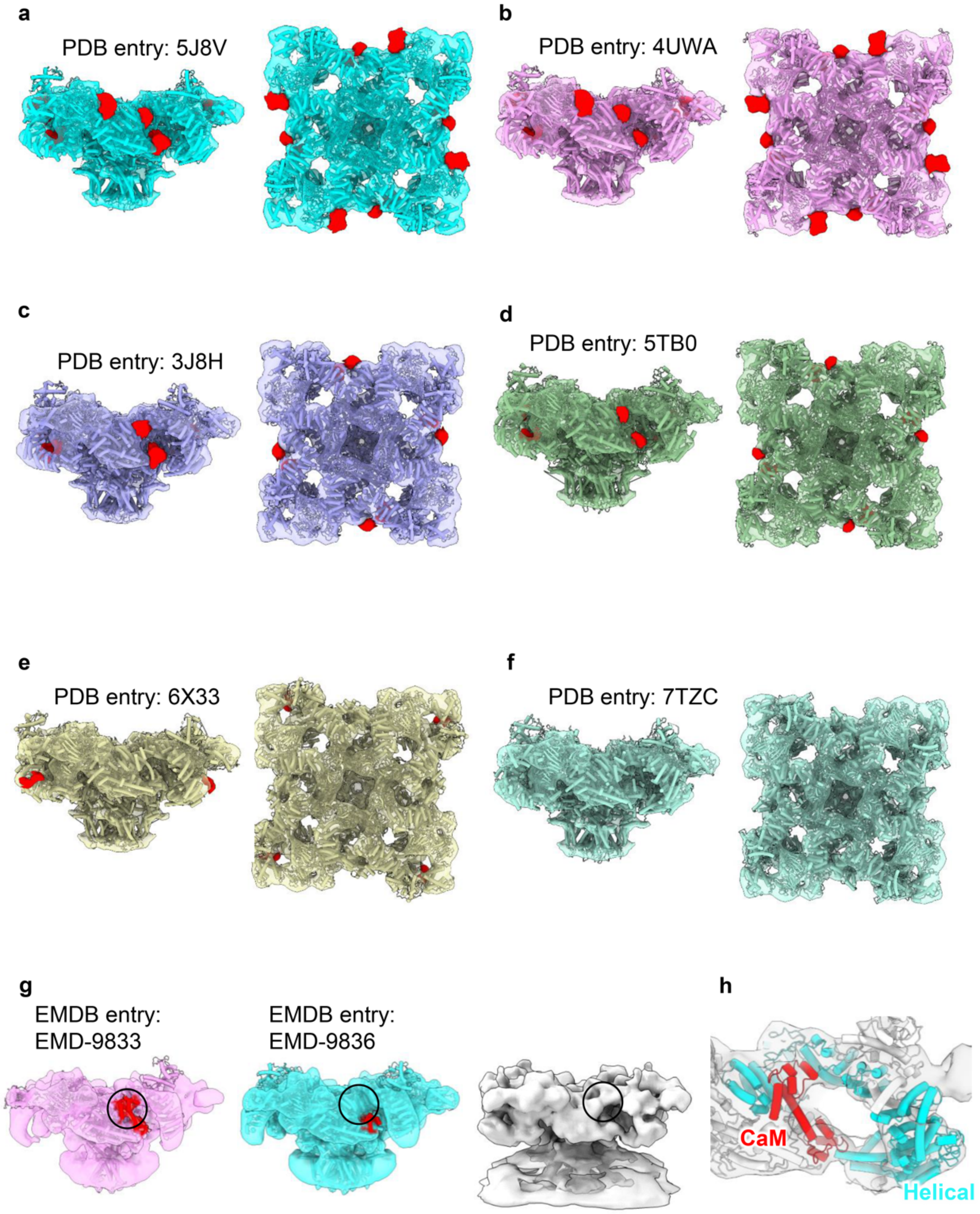


**Figure S3. Comparisons between the *in situ* RyR1 structure and the previously reported structures of purified RyRs.** (**a-f**) The cryo-EM map of *in situ* RyR1 is fitted with the previously solved structures of RyR1s in open state (a, PDB entry 5J8V) (*22*), closed state (b, PDB entry 4UWA) (*18*), FKBP12 bound states (c and d, PDB entries 3J8H and 5TB0) (*19, 21*), and apo-CaM bound states (e and f, PDB entries 6X33 and 7TZC) (*56, 57*), respectively. The different unoccupied pieces of densities in the cryo-

EM map are colored in red. (**g**) The low-pass filtered cryo-EM maps of RyR2-apo-CaM (EMDB entry EMD-9833) (*58*), RyR2- $Ca^{2+}$-CaM (EMDB entry EMD-9836) (*58*), and *in situ* RyR1 (this study) are presented and colored in pink, cyan and grey, respectively. Both the densities of apo-CaM and $Ca^{2+}$-CaM are colored in red. The density missing in RyR2-$Ca^{2+}$-CaM but appearing in other two maps is indicated by the black circles. (**h**) The fitting of apo-CaM model into the density of *in situ* RyR1 cryo-EM map near HD1 and HD2 domains.

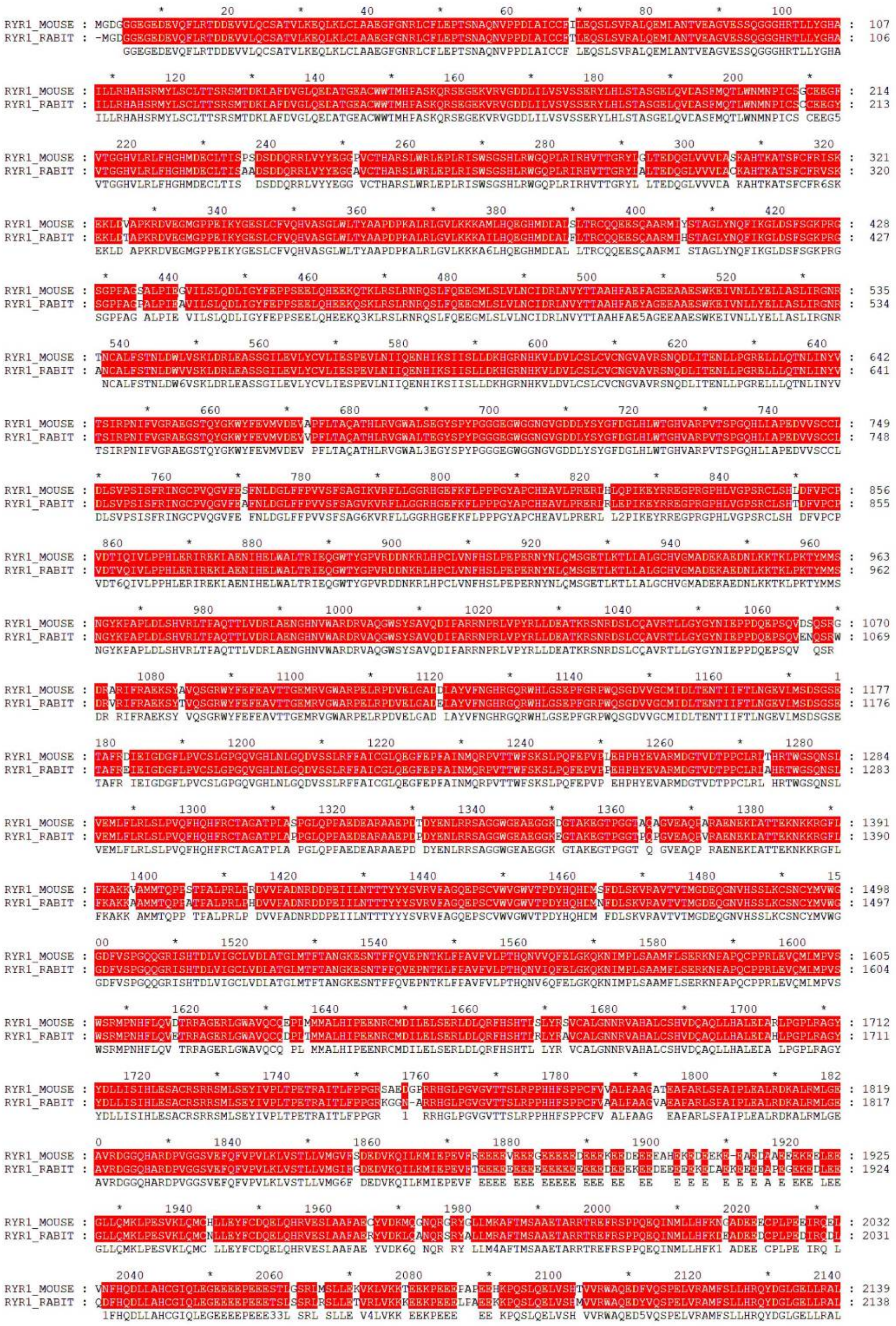


1

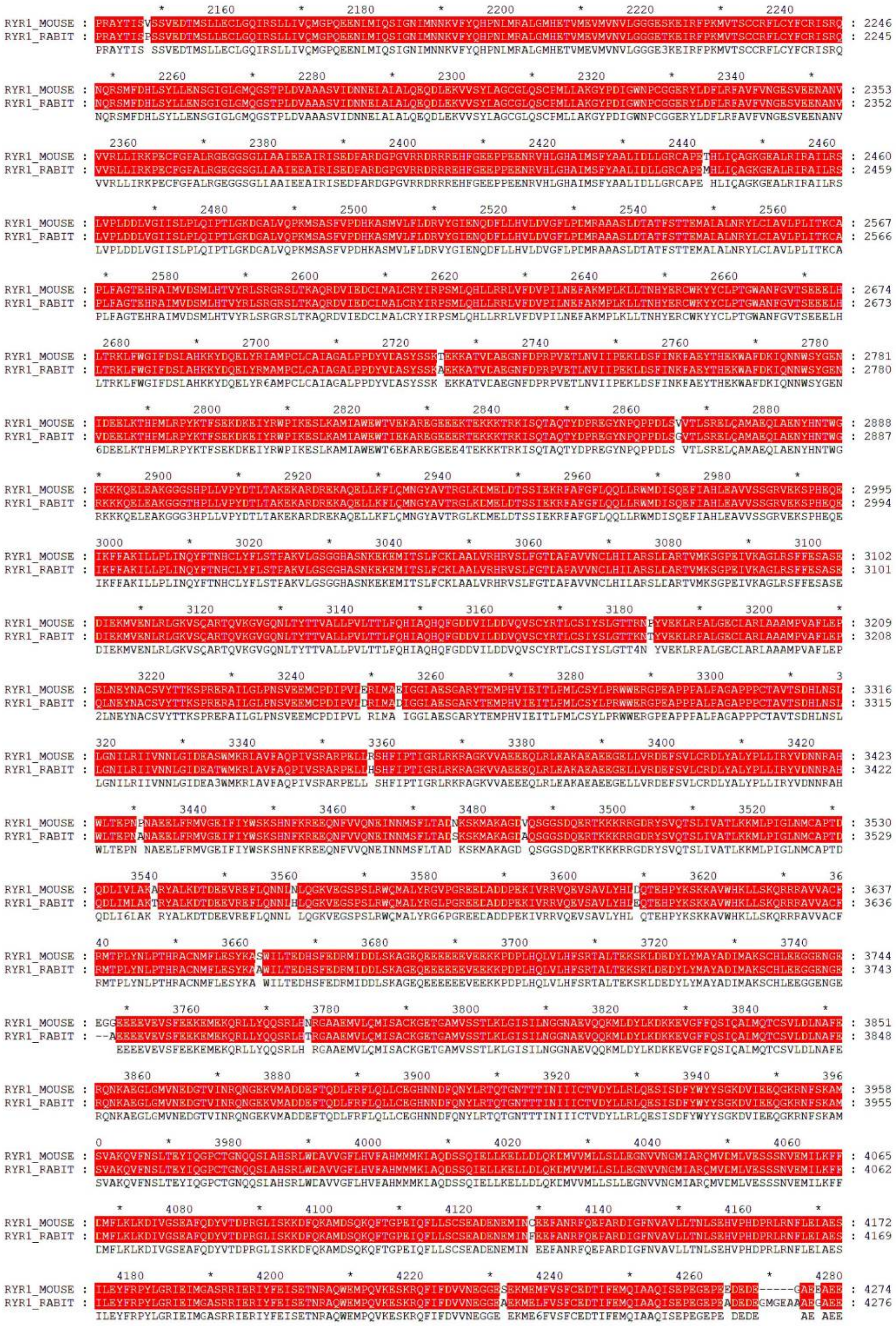

1

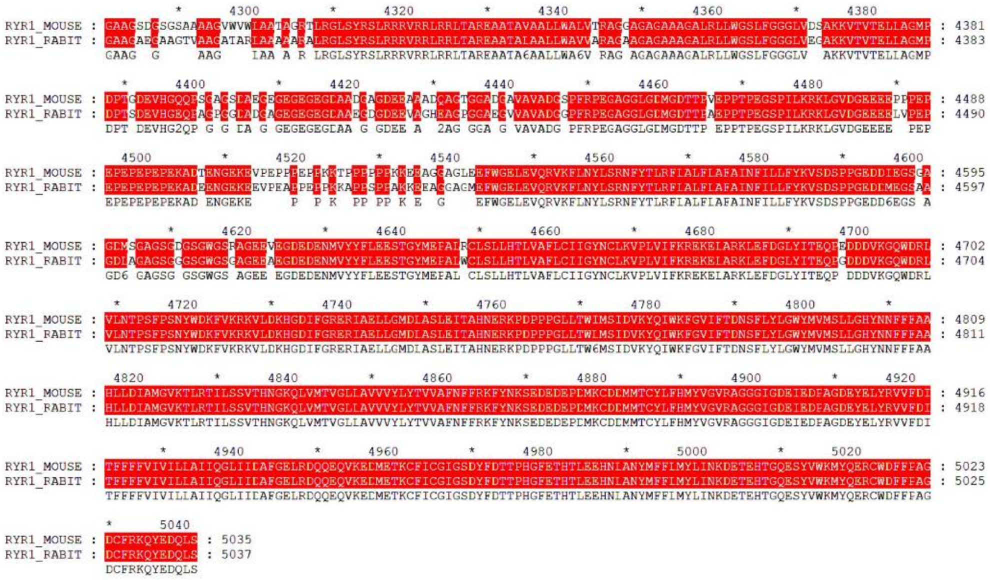

**Figure S4. Sequence alignment of RyR1 sequences between mouse and rabbit.**

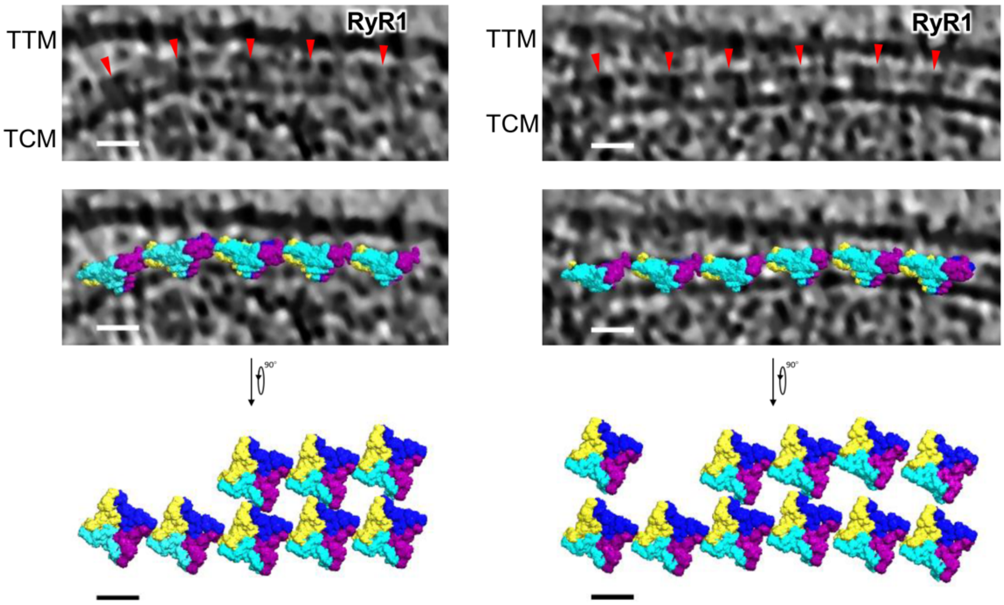


**Figure S5. *In situ* arrangement of RyR1s in native triad junctions.** Another two representative tomograms showing the embedded array of RyR1 tetramers in the triad junctions. The structures of RyR1s are plotted back into the tomograms according to their refined coordinates and Euler angles. Scale bar, 20 nm.

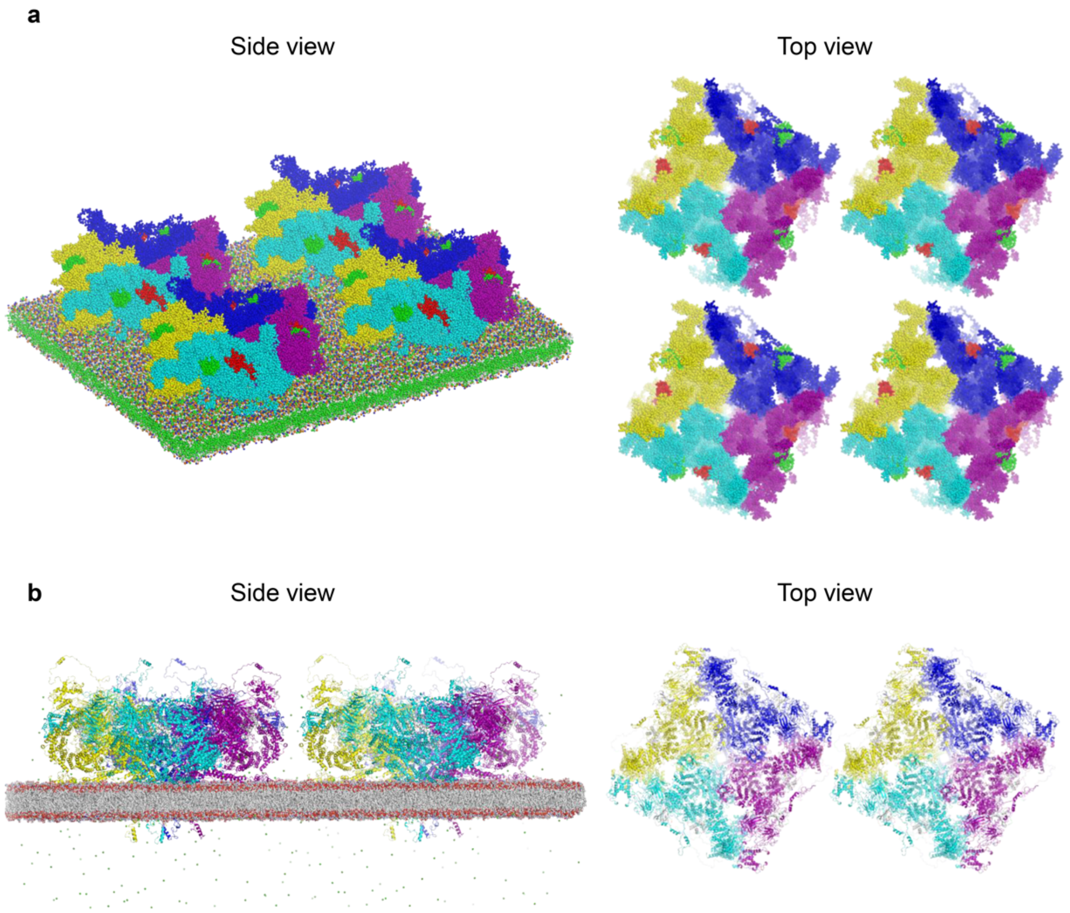


**Figure S6. Initial configurations of molecular dynamics simulations of RyR1 tetramers.** (**a**) Coarse grain MD simulation of four RyR1 tetramers. (**b**) All-atom MD simulation of two RyR1 tetramers. Each RyR1 protomer is complexed with FKBP12 and apo-CaM. The RyR1 tetramers were initially placed larger than 20 Å away from each other. The initial setup systems were similar for both open and closed states.

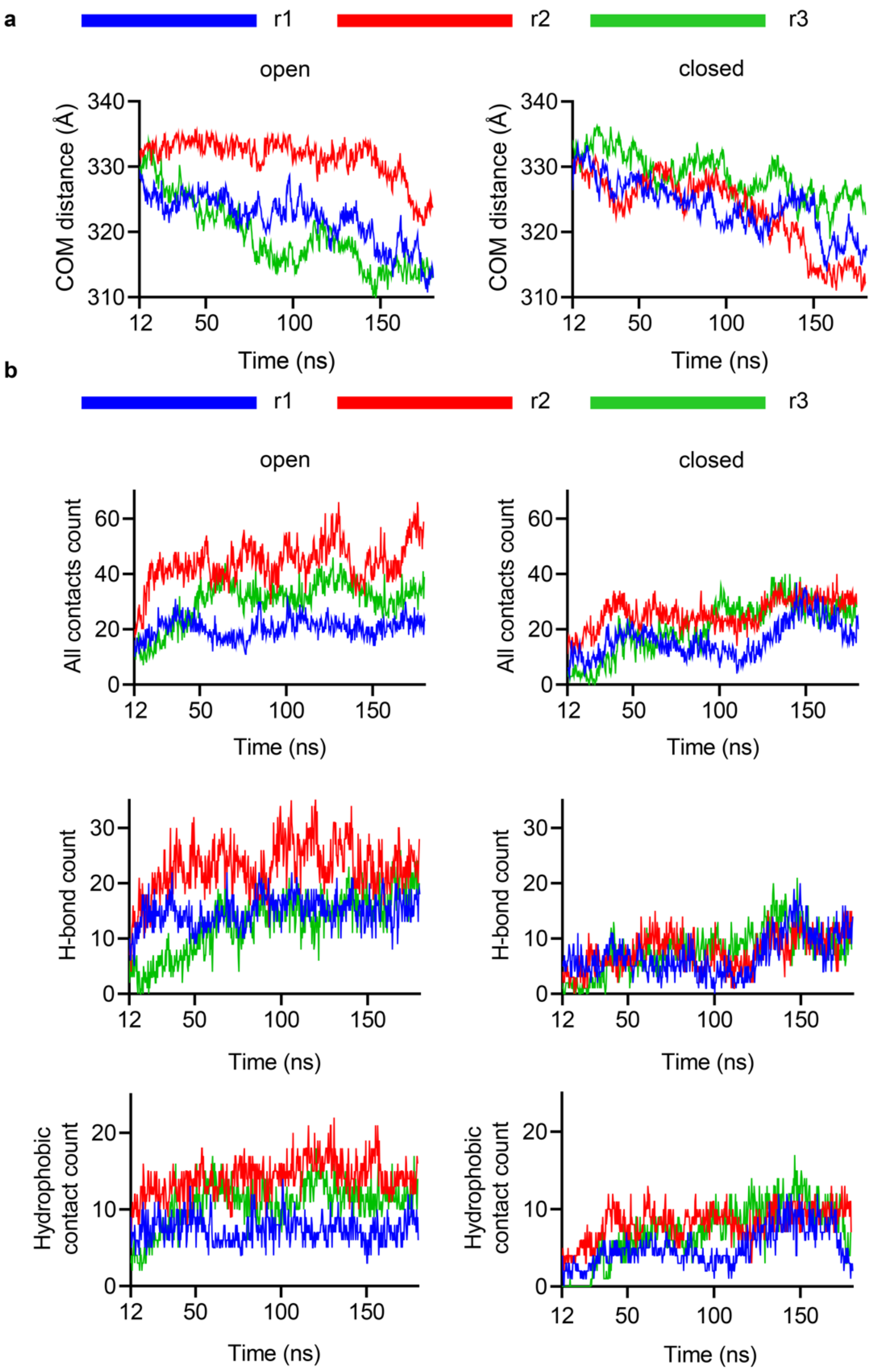

a
r1
r2
r3
open
closed
COM distance (Å)
340
330
320
310
12
50
100
150
Time (ns)
b
All contacts count
60
40
20
0
H-bond count
30
Hydrophobic contact count


1

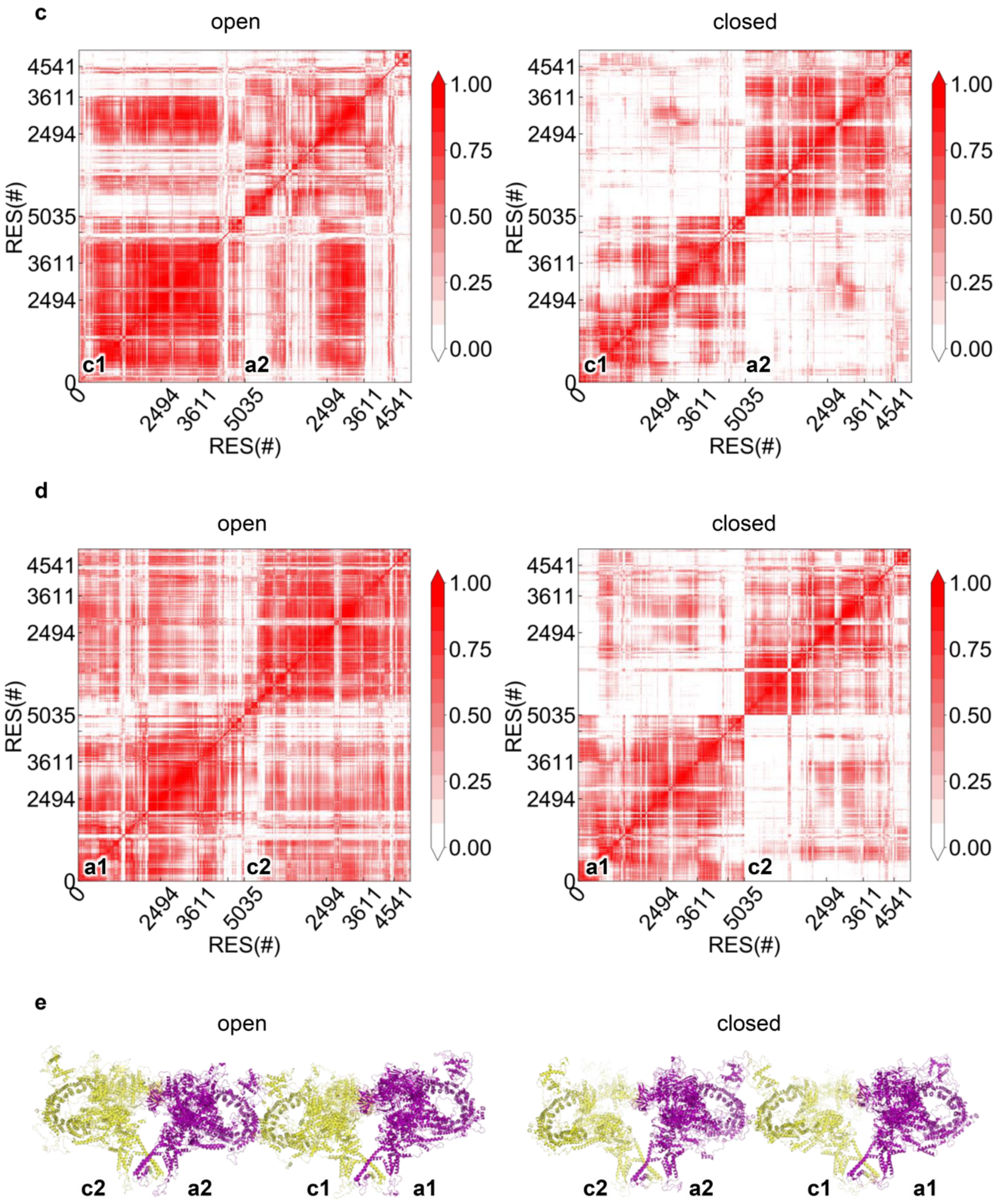


**Figure S7. All-atom MD simulations of RyR1-RyR1 interactions and dynamics.** (**a**) Time evolution profiles of COM distances between two RyR1 tetramers (as illustrated in Fig. 4c) in the open and closed states, respectively. The data was acquired from three MD replicas, labeled as r1, r2, and r3, respectively. (**b**) Time evolution profiles of counts of total contacts (< 6 Å), H-bonds, and hydrophobic contacts (< 6 Å) of two interacting RyR1 tetramers in the open and closed states, respectively. The data was acquired from three MD replicas, labeled as r1, r2, and r3, respectively. (**c**) DCCMs of the interacting

RyR1 protomers in the open and closed states, respectively, which are the full profiles of Fig. 4f. Correlated values above zero are represented in gradient red and the high positive value represents a strong correlated motion. (**d**) DCCMs of two most distant RyR1 protomers, which are labeled as a1 and c2 in (e), in the open and closed states respectively.

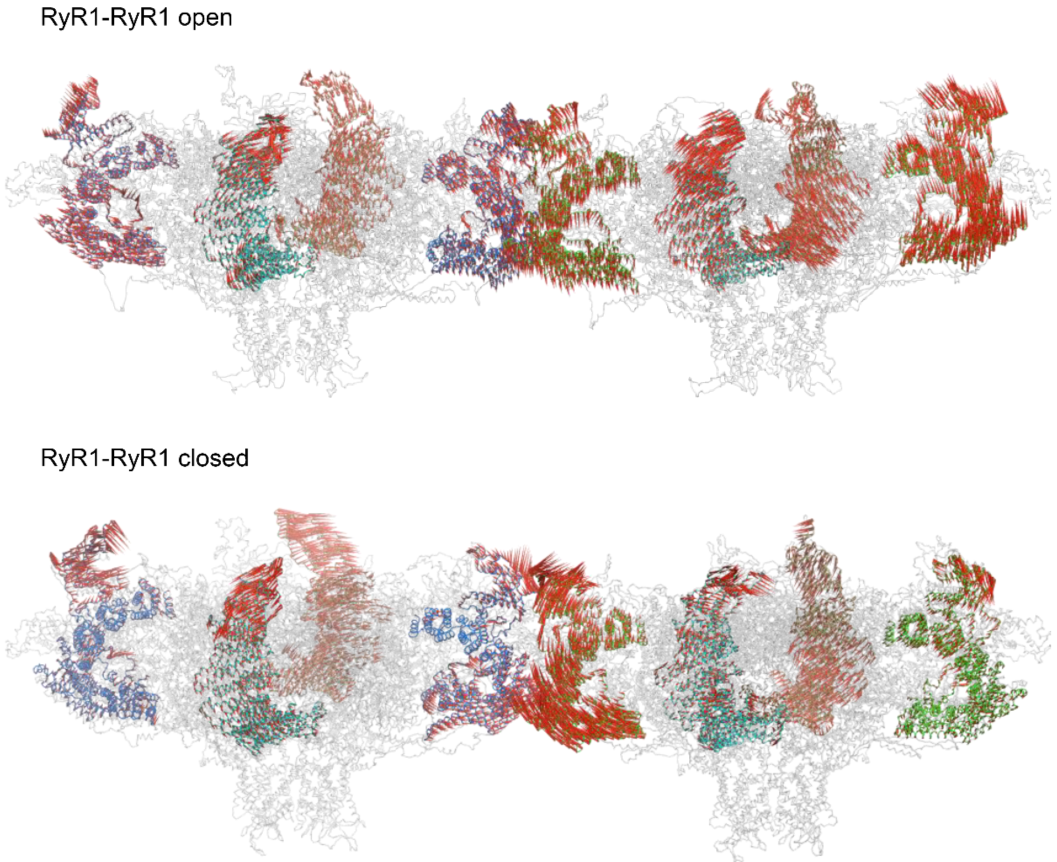


**Figure S8. Synergetic motions of open RyR1s observed from all-atom MD simulations.** The primary motions of RyR1s in both open and closed states were calculated based on the first principal component (PC1s) from PCA investigation of all-atom MD simulations of dimeric RyR1 tetramers. The regions with residues ranging from 2494 to 3611 are colored and the red arrows indicate their motions.

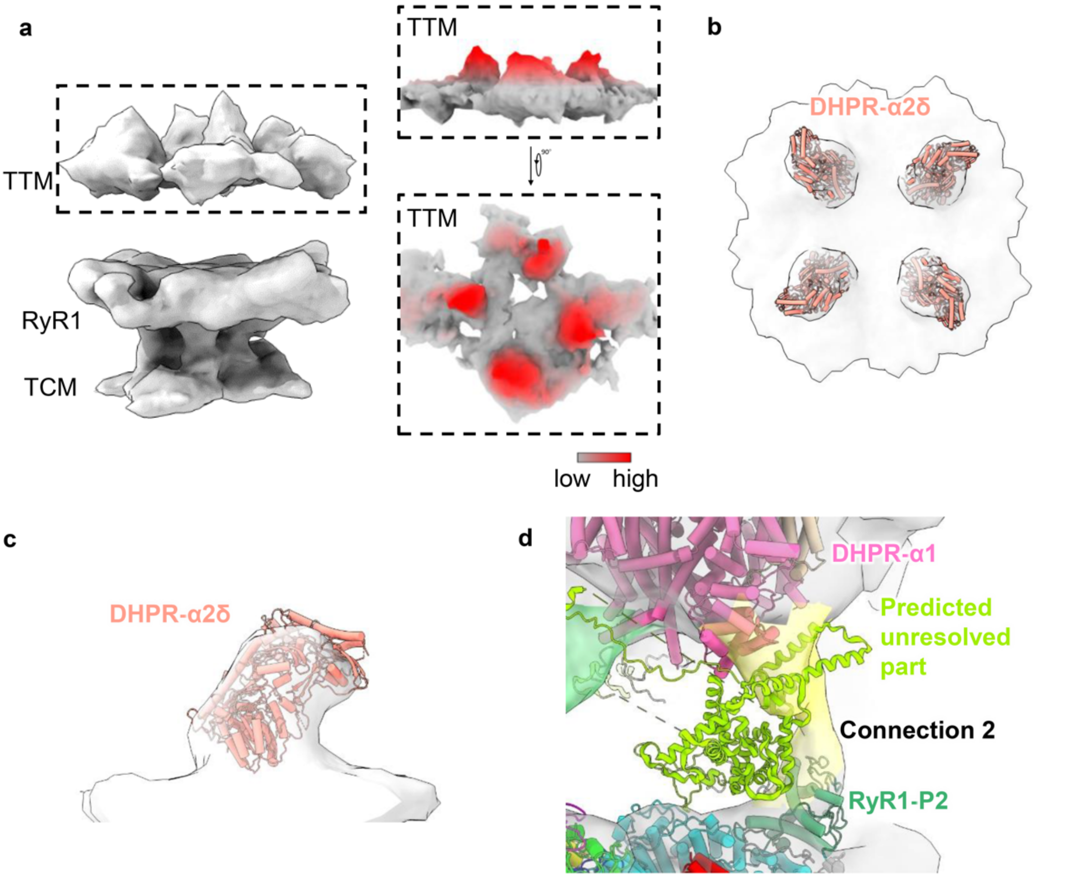


**Figure S9. The structure of RyR1-DHPR supercomplex.** (**a**) The STA averaged map of RyR1-DHPR supercomplex aligned using C1 symmetry. The density of TTM region is shown aside with both side and top views and colored according to the height in the Z-axis direction with the red indicating higher position and grey for lower position. (**b**) and (**c**) Top and side views of extruding parts of RyR1-DHPR supercomplex, which was aligned using C4 symmetry and fitted with four α2δ domains of DHPR (PDB entry 5GJW). (**d**) The predicted structure of DHPR α1 subunit is situated near the density of Connection 2.

### Supplementary Movies

**Movie S1. Representative tomogram of native triad junction.**

**Movie S2. Overall *in situ* structure of RyR1.**

**Movie S3. Arrangement of RyR1s in the native triad junction.**

**Movie S4. A trajectory movie of coarse grained MD simulation of four RyR1 tetramers in the open state.**

**Movie S5. A trajectory movie of coarse grained MD simulation of four RyR1 tetramers in the closed state.**

**Movie S6. A trajectory movie of all-atom MD simulation of two neighboring RyR1 tetramers in the open state.**

**Movie S7. A trajectory movie of all-atom MD simulation of two neighboring RyR1 tetramers in the closed state.**

**Movie S8. Arrangement of RyR1-DHPR supercomplexes in the native triad junction.**

### Supplementary Tables

**Table S1. Statistics of cryo-ET data collection, image processing and model building.**

| | | | | |
|---|---|---|---|---|
| **Data acquisition** | | | | |
| Microscope | Titan Krios G2 | | | |
| Voltage (kV) | 300 | | | |
| Detector | Gatan K2 | | | |
| Energy filter | Gatan GIF Quantum, 20 eV | | | |
| Mode | Gain Normalized | | | |
| Pixel size (Å) | 2.22 | | | |
| Stage tilting angle | About -48° - 32° or -32° - 48° | | | |
| Number of images | 35 - 40 | | | |
| Exposure per tilt (e/Å$^2$) | 4 | | | |
| Exposure per tilt set (e/Å$^2$) | 140 to 160 | | | |
| Defocus range (μm) | -4 - -5 | | | |
| Software | SerialEM | | | |
| **Sub tomogram analysis** | | | | |
| Data set | RyR1 (C4) | RyR1-DHPR supercomplex (C4) | RyR1-DHPR supercomplex (C1) | Dimer of RyR1 tetramer (C2) |
| EMDB ID entry | EMD-37089 | EMD-37092 | EMD-37093 | EMD-37094 |
| Number of tomograms | 230 | 94 | 135 | 227 |
| Number of particles | 1365 | 154 | 251 | 1239 |
| Symmetry | C4 | C4 | C1 | C2 |
| Resolution (Å) | 16.7 | 33.0 | 39.0 | 25.0 |
| Map pixel size (Å) | 4.44 | 8.88 | 8.88 | 8.88 |

**Table S2. Sequence information and templates used for homology modelling during MD simulations.**

| | Residues | Template (PDB entry) |
|---|---|---|
| RyR1 open | 1-5053 | 7CF9 |
| RyR1 closed | 1-5053 | 7M6A |
| FKBP12 | 2-108 | 7CF9 |
| CaM | 5-148 | 6M2W |

**Table S3. A summary of major simulations.**

| Systems | No. of atoms | water No. | Box size (Å$^3$) | MD Simulation (ns) |
|---|---|---|---|---|
| All-atom models | | | | |
| RyR1-open | 3070624 | 770739 | 342×342×282 | 100×3 copies |
| RyR1-closed | 2814268 | 676952 | 344×344×232 | 100×3 copies |
| RyR1-open dimer | 6141248 | 1541478 | 688×344×280 | 180×3 copies |
| RyR1-closed dimer | 5628536 | 1353904 | 685×343×253 | 180×3 copies |
| CG models | | | | |
| RyR1-open 4mer | 1174868 | 198185 | 708×708×255 | 3000×10 copies |
| RyR1-closed 4mer | 1228644 | 211491 | 717×717×262 | 3000×10 copies |
